# Multi-scale biophysical modeling of fungal colony growth

**DOI:** 10.64898/2026.11.05.764668

**Authors:** Lauren Yan, Gianna L. Marschmann

## Abstract

Fungal colony growth is shaped by the fundamental physical operating principles of hyphal elongation, branching, and anastomosis, which collectively organize into macroscopic foraging networks. While sophisticated 3D models utilizing adaptive mesh refinement (AMR) on high-performance computing architectures can explicitly capture these discrete cellular mechanics, their immense computational cost prevents their integration into predictive ecosystem simulators. Here, building upon the Biological Modeling and interface eXchange (BMX) framework, we combine high-fidelity discrete hyphal mechanics with an analytically tractable, stochastic traveling-wave surrogate (sBARE) to provide multiscale simulations of fungal colonies at a fraction of the computational cost. By leveraging demographic fluctuations at the advancing front, our framework uniquely resolves macroscopic parameter degeneracies, separating the directed forward drift of leading tips from their effective diffusion to capture the macroscopic signature of negative autotropism in 3D networks. Simulations reproduce experimentally observed traveling wave profiles and confirm that interior network density under replete conditions emerges as an invariant self-regulated by fusion. Furthermore, global sensitivity analysis reveals that expansion speed and network density are governed by largely disjoint microscopic parameters. We demonstrate that because the physics of hyphal growth permit networks that are both exceptionally fast and dense, the empirical trade-off between sparse ‘explorer’ and dense ‘exploiter’ strategies observed in nature is not a strict mechanical constraint, pointing instead to metabolic limits on local growth rules. Our mathematically tractable approach bridges exascale-level cellular mechanics with macroscopic wave dynamics, enabling the integration of trait-driven advective fluxes and carrying capacities into the spatial transport and reaction network schemes of large-scale biogeochemical models.

**Author summary:** Fungi forage by extending microscopic tubular cells called hyphae, which branch and fuse to form vast networks. A major challenge in biophysics is understanding how these localized rules coordinate to drive a colony expanding like a traveling wave. While supercomputers can simulate these cellular mechanics in high detail, their massive computational cost prevents their use at larger scales. To bridge this gap, we developed a mathematical framework that translates these heavyweight simulations into fast-running, simplified reaction-diffusion equations. By analyzing the “noisy” random wandering of the growing edge, we successfully separated the directed forward growth of tips from their random exploratory movement. Our simulations confirmed that, rather than passively running out of resources, fungi actively control their own network density through fusion, acting as an intrinsic biological population cap. Furthermore, we found that a colony’s expansion speed and its interior density are controlled by completely different cellular traits. Because physical rules allow fungi to theoretically be both fast-growing and dense, the trade-offs we observe in nature—where fungi are typically forced to be either sparse “explorers” or dense “exploiters”—must be driven by metabolic energy limits rather than mechanical constraints. Ultimately, our equations provide a mathematically simple way to incorporate realistic, trait-driven fungal growth into predictive ecosystem simulators.

## Introduction

Understanding how cellular growth rules coordinate the expansion and architecture of biological networks is a fundamental challenge across biophysics and ecological modeling [1–3]. Filamentous fungi represent a striking example of such indeterminate self-organization: lacking a centralized developmental blueprint, a colony expands through the continuous elongation of discrete hyphal segments [4], physical contact and fusion upon collision [5], mechanical contact with the growth surface [6], and stochastic fluctuations in discrete branching and fusion events [7]. Fungal colonies growing on flat surfaces provide an ideal, multiscale model system to investigate how these local biological and physical interactions self-assemble into an interconnected, adaptive foraging network known as a mycelium [8].

Mycelial expansion relies on two complementary processes operating across disparate spatial scales. Apical extension drives branching outgrowths that push the fungal colony outward into unexploited territory [9, 10], while anastomosis reconnects filaments to establish the cytoplasmic continuity required for inter-segment translocation and long-distance nutrient transport [11, 12]. Because fungal growth is indeterminate, understanding mycelial foraging requires linking physical processes across four orders of magnitude: branching and fusion originate from sub-micron vesicle and signaling mechanics within hyphae only a few micrometers wide, yet collective cell dynamics organize an interconnected network spanning centimeters [13].

Recent advances in live-cell imaging have begun to bridge these disparate scales, linking single-hypha biophysics to colony-scale behavior. By combining micron-scale morphological measurements (such as segment lengths and branching angles) with centimeter-scale network topology [14], quantitative imaging across diverse species reveals how local growth rules translate into macroscopic patterns [15–17]. In open planar environments, hyphal growth manifests as a pulled traveling wave [18]: a localized front of active, branching hyphal tips advances at constant speed into uncolonized territory, trailing an expanding network of interconnected filaments behind it. Rather than accumulating to an environmental carrying capacity, the interior biomass density of the trailing network stabilizes at a steady plateau self-regulated by anastomosis [16].

To interpret these dynamics, reaction–diffusion equations have been used to model how mycelial traveling waves propagate across space [19, 20]. Although continuum formulations can capture branching modes and interior steady states through phenomenological kinetic terms [19], they coarse-grain discrete hyphae into continuous density fields, bypassing the microscopic scale altogether. In doing so, these mean-field approximations discard physical processes that operate at the single-hypha level: explicit branching geometry [7], the topological memory that determines whether approaching hyphae collide and fuse [21], and the demographic fluctuations inherent to sparse tip populations at the expanding colony margin [22].

Predicting how macroscopic traveling-wave dynamics emerge from hyphal biophysics therefore requires an explicit representation of local cellular interactions.

In contrast, agent-based models explicitly track individual hyphae, resolving the discrete mechanical contact, branching, fusion, and intercellular transport that govern network self-organization [23]. To capture these dynamics in three-dimensional environments, we build on the Biological Modeling and interface eXchange (BMX) framework [24], an open-source, GPU-accelerated simulation suite built on the AMReX adaptive mesh refinement library [25]. BMX models filamentous fungi as off-lattice, bonded cylindrical segments subjected to low-Reynolds-number hydrodynamic drag, soft-core contact repulsions, and short-range attractive forces. The framework couples these Lagrangian hyphal mechanics to Eulerian continuum grids that resolve external nutrient diffusion and uptake, while simultaneously modeling inter-segment metabolite translocation across septal pores alongside stochastic tip splitting and anastomosis. Yet, even with high-performance GPU acceleration, this microscopic fidelity incurs steep computational costs: simulation time scales rapidly with hyphal segment count [24], rendering discrete simulations intractable for the thousands of evaluations required by global sensitivity analysis, parameter calibration, and ecosystem upscaling. Furthermore, lacking closed-form analytical solutions, agent-based models make it difficult to identify the mathematical drivers of macroscopic colony behavior or evaluate parameter sensitivities directly [26].

Surrogate models offer a computationally tractable alternative, yet existing frameworks navigate a trade-off between fidelity and interpretability. While data-driven machine learning emulators can rapidly approximate simulation outputs [27, 28], they often obscure the biophysical mechanisms linking microscopic hyphal rules to macroscopic colony dynamics [29]. Mechanistic continuum surrogates provide a physically grounded alternative [30]: by parameterizing continuum wave equations directly from agent-based simulations and incorporating stochastic fluctuations from discrete tip events, the surrogate retains the analytical tractability and scalability of continuum physics while preserving the demographic noise of individual hyphae [31]. Such a mapping directly connects colony-scale expansion to underlying cellular mechanics and growth parameters.

Here, we present a multiscale framework that bridges single-hypha growth rules [5, 6, 10] with stochastic traveling-wave theory [32–34] to resolve the biophysical basis of mycelial foraging strategies. Using the agent-based BMX model [24], we simulate discrete elongation, branching, and anastomosis across three-dimensional hyphal networks and couple them to an analytically tractable continuum surrogate, the Branching and Annihilating Range Expansion (BARE) model [16]. By extending multi-objective calibration and profile-likelihood methods [31, 35], we construct continuous response surfaces that map microscopic hyphal traits to macroscopic wave speed, peak tip density, and interior filament saturation. This surrogate mapping reduces global parameter exploration from thousands of GPU-hours to seconds without sacrificing mechanistic interpretability. Furthermore, incorporating demographic noise into the continuum equations resolves the stochastic front wandering of leading tips, providing an independent physical observable that uniquely separates directed forward drift from effective tip diffusion.

Evaluating the calibrated framework yields several fundamental biophysical insights. First, the continuum surrogate confirms that under nutrient-replete conditions, interior filament saturation density is an emergent invariant set by the ratio of tip branching to anastomosis rather than an externally imposed carrying capacity [16]. Second, colony expansion operates as a pulled wave [18] driven by the leading subpopulation of pioneer tips, whose reduced effective diffusivity quantitatively captures the negative autotropism observed in exploring hyphae [36, 37]. Third, global sensitivity analysis reveals that wave speed and saturation density are governed by largely disjoint parameter subsets: wave velocity is driven primarily by apical tip splitting, whereas interior filament density is determined by higher order interactions between segment length, hyphal caliber, and fusion radius. This separation demonstrates that the inverse speed–density relationship is not an unavoidable mechanical constraint [38, 39], indicating that the continuum from fast, sparse explorers to slow, dense exploiters [14, 40] emerges instead from metabolic limits on local growth rules. Together, these results demonstrate that an analytically tractable, stochastic continuum surrogate can recover macroscopic mycelial growth strategies from discrete hyphal biophysics, establishing an operational path to embed mechanistic fungal traits into large-scale biogeochemical models.

## Materials and Methods

### Discrete agent-based simulation of hyphal networks (BMX)

To simulate the three-dimensional growth of filamentous fungi on flat agar surfaces, we used the open-source, high-performance BMX (Biological Modelling and interface eXchange) package — an extension of the AMReX adaptive mesh refinement framework [23, 25]. BMX represents hyphae as off-lattice cylindrical particles and provides particle evolution methods that both transport particles through the simulation domain and mediate two-way coupling between the particles and the underlying adaptively refined continuum grid, allowing hyphae to consume and release metabolites locally while responding to the resulting concentration fields. A mechanical model of anisotropic pairwise interactions controls the forces and bonds between neighboring hyphae and between hyphae and the agar surface.

We varied seven parameters in the BMX model governing hyphal morphology, branching, and fusion (Table 1). The maximum hyphal length *l*_*max*_ and maximum hyphal radius *r*_*max*_ together define the size limits of an individual segment, beyond which volumetric growth ceases. When a tip reaches *l*_*max*_, it undergoes a septation event in which the parent segment is shortened and the excess mass is redistributed into one or two newly created daughter segments; the post-split cylinder length *l*_*split*_ sets the length of these daughter segments, with the parent shortened correspondingly to conserve mass. To ensure that *l*_*split*_ is less than *l*_*max*_, we independently varied the length *dl* = *l*_*max*_*− l*_*split*_. The splitting probability *p*_*s*_ governs the stochastic choice made at each septation event between producing a single new growth tip (continued linear extension) and splitting into two new growth tips (apical branching), and is therefore the sole determinant of branching density in the model, since lateral branching is not currently supported. Anastomosis is restricted to tip-to-interior fusion: at every time step, the distance from each growth tip to every interior segment is checked, and if this distance falls below the maximum fusion separation distance *r*_*f*_, a fusion event occurs with probability *p*_*f*_ . When fusion occurs, a septum is introduced in the target segment at the point of contact, dividing it into two smaller segments and establishing cytoplasmic continuity between the fusing tip and the rest of the network. Finally, the width of the attractive region *W*_*A*_ sets the range of the short-range attractive force between approaching hyphae, controlling the distance over which neighboring segments mechanically influence one another before contact.

**Table 1.** Microscopic hyphal parameters varied in the agent-based (BMX) simulation design.

| **Description** | **Parameter** | **Sampling Range** |
|---|---|---|
| Maximum segment length | *l*<sub>*max*</sub> | 17.5 − 52.5 *µm* |
| Post-split segment length difference | *dl* | 2.5 − 7.5 *µm* |
| Maximum segment radius | *r*<sub>*max*</sub> | 1.25 − 3.75 *µm* |
| Tip-splitting probability | *p*<sub>*s*</sub> | 0.1 − 0.3 |
| Fusion probability per time step | *p*<sub>*f*</sub> | 0.005 − 0.015 |
| Maximum fusion separation | *r*<sub>*f*</sub> | 3 − 9 *µm* |
| Width of hyphal attraction region | *W*<sub>*A*</sub> | 0.5 − 1.5 *µm* |

Seven agent-based model parameters governing hyphal morphology, branching frequency, and anastomosis geometry were varied across a Sobol sequence ( *±* 50% around baseline values). For the complete list of model parameters and fixed values, see [24].

Each simulation was initialized with a single spore and run for 34 simulated hours (500,000 time steps), with the network state saved at one-hour intervals. Input settings for the seven varied parameters were drawn from the BMX parameter space using a Sobol sequence [41], with each parameter sampled over a range of ±50% around its default value. Three replicates were generated for each sample to capture the average behavior and account for stochasticity, yielding 128 unique parameter samples × 3 replicates = 384 total simulations. Simulations were performed on the Einsteinium (Lawrencium) GPU cluster using NVIDIA V100 GPUs (CUDA target architecture 7.0). Each production run used a single V100 with 4 CPU cores and required approximately 4 hours of wall time, for a total of 1,500 GPU-hours.

To analyze BMX output, we developed a post-processing pipeline that extracts segment densities, tip kinematics, and branching and fusion event statistics from the saved particle data. BMX outputs particle-level data in which each segment is labeled as a growth tip, an interior segment, or a secondary segment (the segment directly behind a tip). To compute segment densities, we constructed the convex hull of the network at each saved time step and partitioned the interior into concentric rings of equal width obtained by uniformly scaling the hull about its centroid. The density in each ring was defined as the total segment count divided by the ring area. Using the segment-type labels, we separately computed densities for growth tips and interior filaments. Growth tips located at a vertex of the convex hull were further classified as “puller hyphae”.

To compare dynamics across rings, time was shifted into a ring-local reference frame in which *t*_0_ corresponds to the time at which the ring reaches half its maximum filament density. The value of *t*_0_ was obtained for each ring by fitting the filament density over time to a sigmoid,

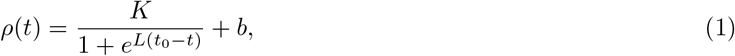

with *K, L, t*_0_ and *b* as free parameters.

Tip velocities **v** were computed as the average displacement of each growth tip between successive time steps. The drift velocity was defined as the component of **v** along the local bulk flow direction, taken to be the outward radial unit vector 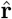 from the center of the final network: *v*_*d*_ = |**v**| cos *θ*, where *θ* is the angle between **v** and 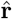.

To track branching and fusion rates, we added a global event counter to BMX that increments whenever a branching or fusion routine completes and records the unique identifier of the segment involved. This was merged with the particle data to determine the location of events. We then partitioned the network into concentric rings about its center and computed, within each ring, the time-averaged tip density *n*, filament density *ρ*, branching rate *b*, and fusion rate *a*. Linear regressions of *b* against *n* and of *a* against *nρ* yield the proportionality constants in the assumed linear forms for the tip splitting and fusion rates.

### Stochastic continuum traveling-wave surrogate model (sBARE)

The Fisher–Kolmogorov (FKPP) equation is commonly used to model population range expansion as a traveling wave [42, 43]. The Branching and Annihilating Range Expansion (BARE) model modifies the FKPP equation to capture the essential branching and fusion dynamics of fungal hyphae— which constitute the expanding front and a population size control— and a distinction between the outer growing tips and the filaments they lay down [16].

BARE coarse-grains the mycelium into two coupled continuous fields on the colony plane **r** ∈ ℝ^2^: the active growing-tip density *n*(**r**, *t*) (mm^−2^), which drives outward front expansion, and the persistent filament density *ρ*(**r**, *t*) (mm^−1^), representing the total length of hyphae deposited per unit area. The separation into a mobile, reactive tip field and an immobile, accumulating filament field encodes the central biological fact that only tips extend, branch, and fuse, whereas the filament they lay down is essentially static on the timescale of front propagation.

The spatial transport of growing tips is governed by an advection–diffusion constitutive relation,

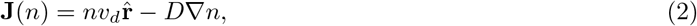

where the drift velocity *v*_*d*_ (mm hr^−1^) captures the outward directional persistence of tips along the radial propagation vector 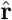, and the effective diffusivity *D* (mm^2^ hr^−1^) coarse-grains the random reorientation and angular dispersion introduced by stochastic branching events.

The deterministic two-field BARE system is:

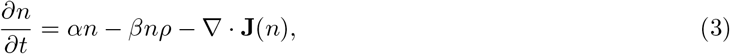

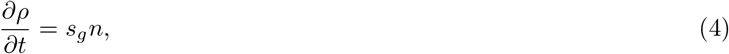

where *α* (hr^−1^) is the per-capita branching rate constant, *β* (mm hr^−1^) is the anastomosis rate constant, and *s*_*g*_ (mm hr^−1^) is the mean elongation rate. The model has five parameters (Table 2), **x** = (*α, β, v*_*d*_, *s*_*g*_, *D*), which we infer by matching to BMX output.

**Table 2.**
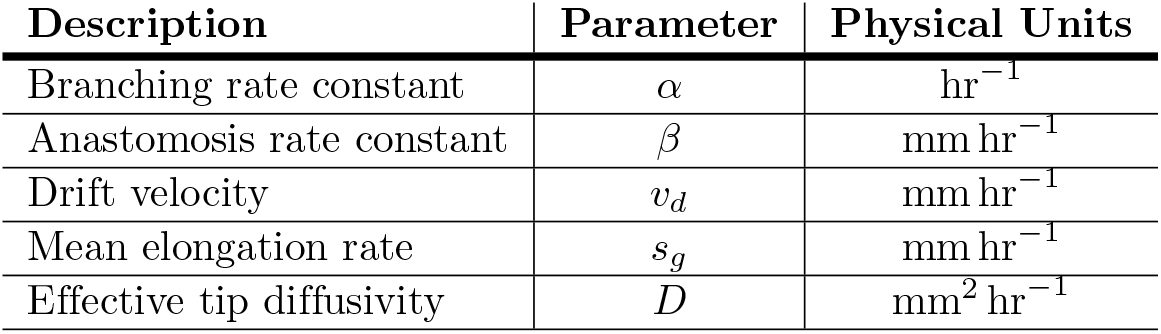
Macroscopic parameters of the BARE traveling-wave continuum model.

| **Description** | **Parameter** | **Physical Units** |
|---|---|---|
| Branching rate constant | *α* | hr<sup>−1</sup> |
| Anastomosis rate constant | *β* | mm hr<sup>−1</sup> |
| Drift velocity | *v<sub>d</sub>* | mm hr<sup>−1</sup> |
| Mean elongation rate | *s<sub>g</sub>* | mm hr<sup>−1</sup> |
| Effective tip diffusivity | *D* | mm<sup>2</sup> hr<sup>−1</sup> |

Macroscopic parameters governing tip kinetics, spatial transport, and filament deposition in the Branching and Annihilating Range Expansion (BARE) continuum model [16].

In Eq. (3), branching is modeled as a first-order autocatalytic process (*αn*): each tip branches at a constant per-capita rate, such that the production of new tips is proportional to the local tip density and independent of proximity to other tips. Anastomosis is modeled as a mass-action fusion process between tips and existing filaments (*βnρ*): an active tip is annihilated at a rate proportional to both the local tip density *n* and the surrounding filament density *ρ* available for fusion. The linearity of the branching term in *n* is decisive. Because the per-tip growth rate *α* does not depend on *n* (in particular it does not vanish or diminish at small *n*), the reaction term is maximal in the dilute leading edge. Fronts of this type—where the growth rate is largest where the population is smallest—are pulled: the propagation is controlled entirely by the dynamics of the sparse tips at the leading edge, and a small number of tips there sets the speed of the whole invasion [34]. This classification is what licenses the FKPP marginal-stability speed selection used below, and, later, the finite-*N* corrections of pulled-front theory [18, 43].

In the region behind the leading edge, curvature of the front is negligible over the front width, so we treat propagation locally as planar along the radial coordinate *r* and introduce the co-moving coordinate *ξ* = *r− ct*, with *c* the (as yet unknown) front speed. Under the travelling-wave ansatz *n*(*r, t*) = *n*(*ξ*), *ρ*(*r, t*) = *ρ*(*ξ*), the time derivatives transform as ∂_*t*_(*·*) =−*c* ∂_*ξ*_(*·*), and Eqs. (3)–(4) become the ordinary differential equations

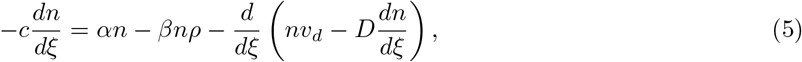

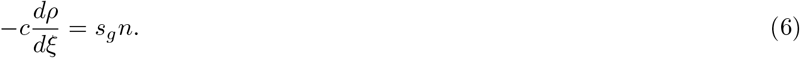

We orient *ξ* so that the uninvaded region (*n* → 0, *ρ* → 0) lies at *ξ* → +∞ and the colonized interior lies at *ξ* →*−*∞.

#### Closed form for the saturation density

To derive the steady-state interior filament density, *ρ*_sat_ ≡ *ρ*(*ξ* →−∞), we integrate the co-moving traveling-wave system across the entire front (*ξ* ∈ [−∞, +∞ ]). Integrating Eq. (6) across the wave profile yields

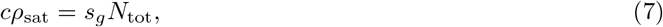

where 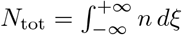 is the total tip count per unit transverse length, and we apply the boundary conditions *ρ*(+ ∞ ) = 0 and *ρ*(−∞) = *ρ*_sat_.

Next, integrating Eq. (5) across the front eliminates the transport terms, as both *n* and its spatial gradient vanish at the boundaries (*n*(*±*∞) = 0, *n*^*′*^(*±*∞) = 0). The steady-state tip balance simplifies to:

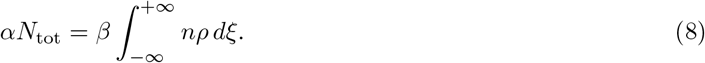

Using the relation *n* =−(*c/s*_*g*_) *dρ/dξ*, the non-linear tip–filament collision integral evaluates directly:

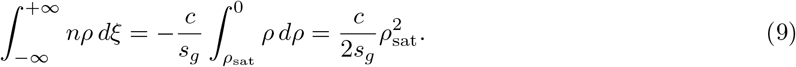

Substituting Eqs. (7) and (9) into Eq. (8) yields the exact closed form invariant:

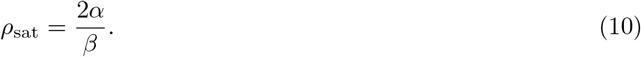

The saturation density is set purely by the ratio of branching to anastomosis and is independent of transport (*v*_*d*_, *D*) and of the elongation rate *s*_*g*_. It provides a clean, transport-free constraint on *α/β*.

#### Closed form for the front speed

Because the wave is pulled, its speed is fixed by the linearized dynamics at the leading edge, where *n* → 0 and *ρ* → 0. Linearizing Eq.(3) about the unstable state (*n, ρ*) = (0, 0) (the anastomosis term *βnρ* is second order in the small fields and drops out) gives the constant-coefficient equation

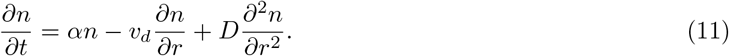

Seeking exponentially decaying leading-edge modes *n*∼ *e*^−*λ*(*r*−*ct*)^ with *λ >* 0 yields the dispersion relation between admissible decay rate *λ* and wave speed *c*,

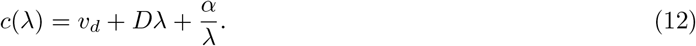

For a pulled front the selected speed is the marginal-stability (minimal) speed over admissible *λ*; minimizing Eq.(12),

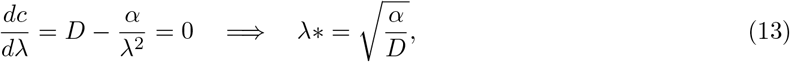

and substituting *λ*^*^ back gives

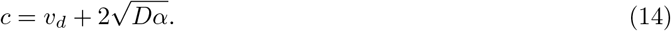

This is the FKPP/marginal-stability speed shifted rigidly by the drift velocity *v*_*d*_, exactly as anticipated by the Galilean argument: in the frame moving at *v*_*d*_ the front advances at the pure FKPP speed 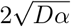.

#### The (*v*_*d*_, *D*) degeneracy

Equation (14) exposes astructural non-identifiability. For fixed *α*, the deterministic speed constrains only the combination 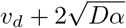: a continuum of (*v*_*d*_, *D*) pairs lying along the curve

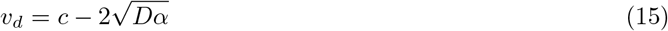

produce identical fronts at the deterministic level. The saturation density (Eq. (10)) is transport-free, and the wave amplitude constrains (*v*_*d*_, *D*) only weakly and non-uniquely. Any fit based solely on mean-field quantities will therefore return a ridge of equally good solutions. Breaking the degeneracy in Eq. (15) thus requires an observable that distinguishes *v*_*d*_ from *D*; front fluctuations supply exactly such an observable.

#### Demographic-noise (stochastic) BARE

We promote Eq. (3) to a stochastic partial differential equation by incorporating demographic noise arising from discrete tip events [22]. For a birth-limited pulled process, the leading fluctuation is multiplicative square-root (Itô) noise:

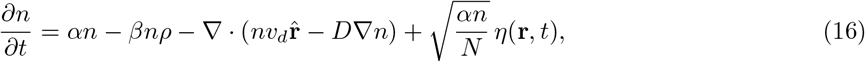

where *η*(**r**, *t*) is Gaussian white noise, ⟨ *η*(**r**, *t*)*η*(**r**^*′*^, *t*^*′*^)⟩ = *δ*(**r− r**^*′*^)*δ*(*t− t*^*′*^), and *N* sets the number of tips per unit reference area, i.e., the coarse-graining scale that controls the amplitude of demographic fluctuations. Eq. (16) corresponds to the stochastic Fisher–KPP (sFKPP) equation with Galilean drift *v*_*d*_, which enters as an outward convection term that does not couple directly to the noise amplitude.

The primary control parameter is not *N* per unit area, but the effective number of tips within the active front region, *N*_*e*_. Defining the characteristic pulled-front scale as 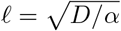, the effective tip number is:]

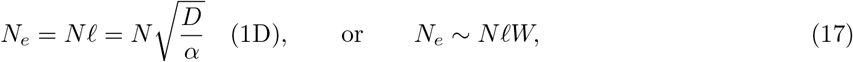

for a front of transverse extent *W* . We measured *N*_*e*_ directly from BMX simulations by counting active tips within one front width of the leading edge; it’s magnitude determines whether finite-*N* corrections below are large enough to be informative.

#### Finite-*N* speed correction

Although fluctuations on the order of 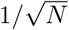 are negligible in dense interior populations, their impact becomes significant at the sparse leading edge (*n* ∼ 1*/N*), where the physical discreteness of individual tips imposes an effective cutoff on the spatial decay of the front. Pulled-front theory demonstrates that this discrete cutoff reduces the selected wave velocity below the deterministic mean-field speed [32–34]. In the co-moving frame translating at drift velocity *v*_*d*_, the leading finite-*N* correction to the wave speed is given by:

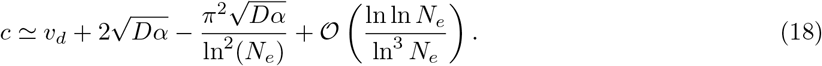

The speed deficit 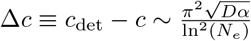 scales with 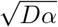 and therefore provides information about *D* independent of *v*_*d*_.

#### Front wandering as a degeneracy-breaking observable

The most informative Galilean-invariant observable is the diffusive wandering of the instantaneous front position, *R*(*t*). While the ensemble mean front advances at constant speed *c*, demographic fluctuations grow diffusively [18, 22, 32],

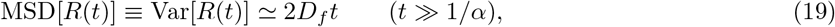

defining the front-wandering diffusivity *D*_*f*_ . Pulled-front theory predicts that *D*_*f*_ follows the asymptotic scaling,

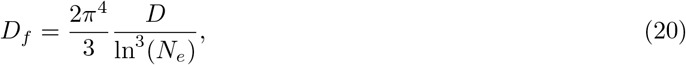

i.e., *D*_*f*_ decays logarithmically as (ln *N*_*e*_)^−3^, with an amplitude governed solely by leading-edge diffusivity *D* and tip count *N*_*e*_, completely independent of directed drift *v*_*d*_.

We determined *D*_*f*_ empirically for each BMX parameter setting by evaluating the mean squared deviation of the hourly wavefront position across *M* stochastic realizations:

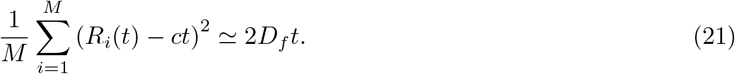

Measuring *D*_*f*_ from discrete BMX simulations uniquely isolates *D* via Eq. (20), after which the directed drift velocity *v*_*d*_ follows directly from the deterministic speed relation 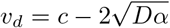, breaking the (*v*_*d*_, *D*)parameter degeneracy.

### Multi-objective calibration and surrogate mapping

We calibrated the continuum BARE model against discrete BMX simulations using three macroscopic quantities of interest:

1. **Wave amplitude:** the peak value of the tip density profile, *n*_max_ = max_*ξ*_ *n*(*ξ*).
2. **Wave front position:** the radial position of the wave peak, whose slope over time defines the wave speed 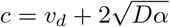.
3. **Saturation density:** the steady-state filament density in the colonized interior, defined analytically by *ρ*_sat_ = 2*α/β*.

For each BMX simulation output, we parameterized BARE by minimizing the weighted squared residuals:

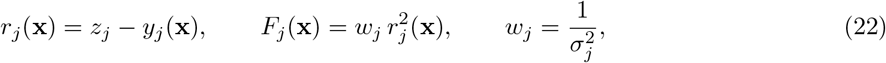

where *z*_*j*_ is the *j*-th macroscopic observable extracted from the BMX spatial wave profile, *y*_*j*_(**x**) is the corresponding BARE solution for parameter vector **x** = (*α, β, v*_*d*_, *s*_*g*_, *D*) in admissible phase space ***χ***, and *σ*_*j*_ is the standard deviation across stochastic BMX realizations.

Because the three observables constrain distinct dynamical features of the traveling wave, we formulated the calibration as a multi-objective optimization problem:

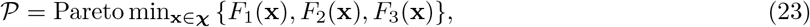

obtaining the set *P* of non-dominated parameter vectors that minimize each residual without degrading the others. We solved this optimization using the AMALGAM evolutionary algorithm [35], which adaptively combines multi-method search operators to converge onto the Pareto front. Initial parameter guesses were obtained from measurements on the BMX output and seeded into the first generation of the evolutionary algorithm. To quantify parameter uncertainty, we fitted the distribution of the final Pareto generation and computed quantiles to establish 95% confidence intervals on the best-fit parameters.

#### Response surface construction and uncertainty quantification

To construct a continuous mapping from discrete hyphal growth rules to macroscopic wave dynamics, we treated each BARE parameter as an implicit function of the microscopic BMX parameter space [31]. For each continuum parameter, we interpolated the upper and lower 95% confidence bounds across the seven-dimensional BMX domain using radial basis function interpolation (*scipy*.*interpolate*.*RBFInterpolator*). At any microscopic input vector 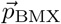, the interpolants return a bounded interval for each continuum parameter. The Cartesian product of these intervals defines a hyperrectangle 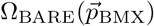 representing the continuum parameter space consistent with that discrete simulation. The macroscopic output metric 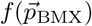 is then evaluated by marginalizing over this parameter mapping uncertainty:

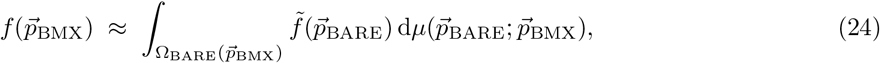

where 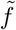 is the continuum model evaluation and *µ* represents a uniform distribution over the hyperrectangle Ω_BARE_. In practice, this integral is approximated by drawing uniform Monte Carlo samples across Ω_BARE_ and computing the ensemble mean.

For surrogate training and cross-validation, input and output dimensions were normalized using min–max scaling. The simulation dataset was partitioned into a 70% training set and a 30% held-out test set. We performed leave-one-out cross-validation (LOOCV) on the training set to optimize the RBF basis kernel, smoothing parameter, and shape factors, and subsequently evaluated predictive accuracy on the unseen test set to confirm that the surrogate generalizes without overfitting. Marginalizing over Ω_BARE_ reduces computationally expensive agent-based evaluations to rapid evaluations of the surrogate, establishing the foundation for global sensitivity analysis.

### Global sensitivity analysis of agent-based model parameters

The calibrated continuum surrogate evaluates in milliseconds while accounting for parameter mapping uncertainty (Eq. (24)). It thus enables global sensitivity analysis across the agent-based model parameter space at negligible computational cost. Let *ρ* denote a probability distribution over the microscopic BMX parameter space representing biological variation across the sampling bounds (Table 1). The global sensitivity of an output observable *f* with respect to the *m* = 7 varied microscopic parameters, GS(*f*(*·*); *ρ*) ∈ *ℝ*^m^, is approximated directly through the surrogate mapping:

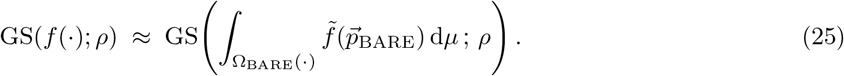

Since global sensitivity methods require only repeated evaluations of the model output together with the input distribution *ρ*, and remain agnostic to the internal numerical implementation of *f*, this formulation allows any standard variance- or distribution-based sensitivity framework to be applied across the agent-based parameter domain. We evaluated the sensitivity of BARE continuum parameters (*α, β, v*_*d*_, *s*_*g*_, *D*) and macroscopic observables (*c, ρ*_sat_) using two complementary frameworks implemented in the *SALib* Python library [44]: variance-based Sobol’ analysis and distribution-based PAWN analysis.

#### Variance-based Sobol’ sensitivity analysis

Sobol’ analysis decomposes the total variance of a model output *Y* = *f*(**X**) into orthogonal contributions from individual input parameters and their higher-order interactions [45]:

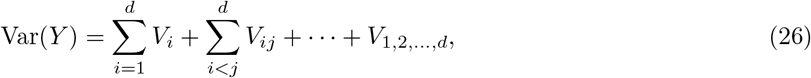

where *d* is the number of parameters, 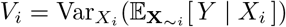 is the first-order variance attributable to parameter *X*_*i*_ alone, and *V*_*ij*_ represents the variance due to second-order interactions between parameters *X*_*i*_ and *X*_*j*_. We evaluated the first-order index, *S*_*i*_ = *V*_*i*_*/*Var(*Y* ), representing the main effect of parameter *X*_*i*_ in isolation, and the total-order index, 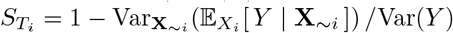, which captures all first-order and higher-order interaction effects involving *X*_*i*_. The total-effect index, i.e., the difference 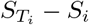 provides a direct diagnostic of non-linear parameter interactions within the surrogate mapping. Quasi-random parameter matrices were generated using Saltelli’s sampling scheme with *N* = 1, 024 base samples, resulting in *N*(2*d* + 2) = 16, 384 surrogate evaluations.

#### Distribution-based PAWN sensitivity analysis

When model output distributions are skewed, multimodal, or heavy-tailed, variance-based decompositions can obscure non-linear parameter influences. PAWN provides a complementary, moment-independent framework that measures parameter sensitivity through shifts in the cumulative distribution function (CDF) of the output [46]. For each input *X*_*i*_, sensitivity is quantified by the Kolmogorov–Smirnov (KS) distance between the unconditional output distribution *F*_*Y*_ (*y*), obtained by varying all inputs simultaneously, and the conditional output distribution 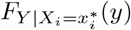, obtained by fixing *X*_*i*_ at a conditioned value 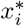 while sampling all other parameters:

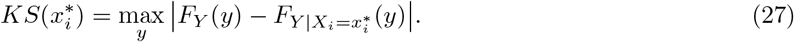

The global PAWN sensitivity index *T*_*i*_ is defined as a summary statistic (e.g., the median) of the KS statistic evaluated across *n*_*c*_ intervals spanning the domain of *X*_*i*_:

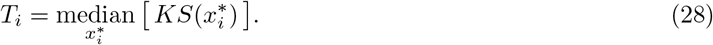

A high PAWN index (*T*_*i*_ → 1) indicates that fixing parameter *X*_*i*_ significantly shifts the entire probability distribution of the macroscopic wave observable, whereas *T*_*i*_ ≈ 0 indicates that the output distribution is insensitive to perturbations in *X*_*i*_. We sampled *N*_uncond_ = 2, 000 unconditional points and evaluated conditional CDFs across *n*_*c*_ = 10 conditioning intervals per parameter. Confidence intervals and standard errors for both Sobol’ and PAWN sensitivity indices were estimated using SALib’s default bootstrap resampling (95% confidence level, [44]).

## Results

### Traveling-wave dynamics emerge from local growth rules

We first asked whether discrete, agent-based BMX simulations reproduce the traveling-wave dynamics characteristic of macroscopic mycelial colony growth [16]. Visualizing the simulated network expansion over time confirms a steadily advancing front of active tips that leaves behind a stationary mesh of hyphal filaments (Fig.1A). To quantify this behavior, we tracked the densities of active growing tips (*n*) and hyphal filaments (*ρ*) across space and time. We found that these spatial profiles maintain a consistent shape as they expand outward (Fig.1B, left panels). This shape-preserving expansion demonstrates the translational symmetries expected of a stable traveling wave: the colony propagates at a constant speed *c* (Fig.1D) and reaches a nearly uniform interior saturation density *ρ*_sat_ (Fig.1C).

**Fig 1.**
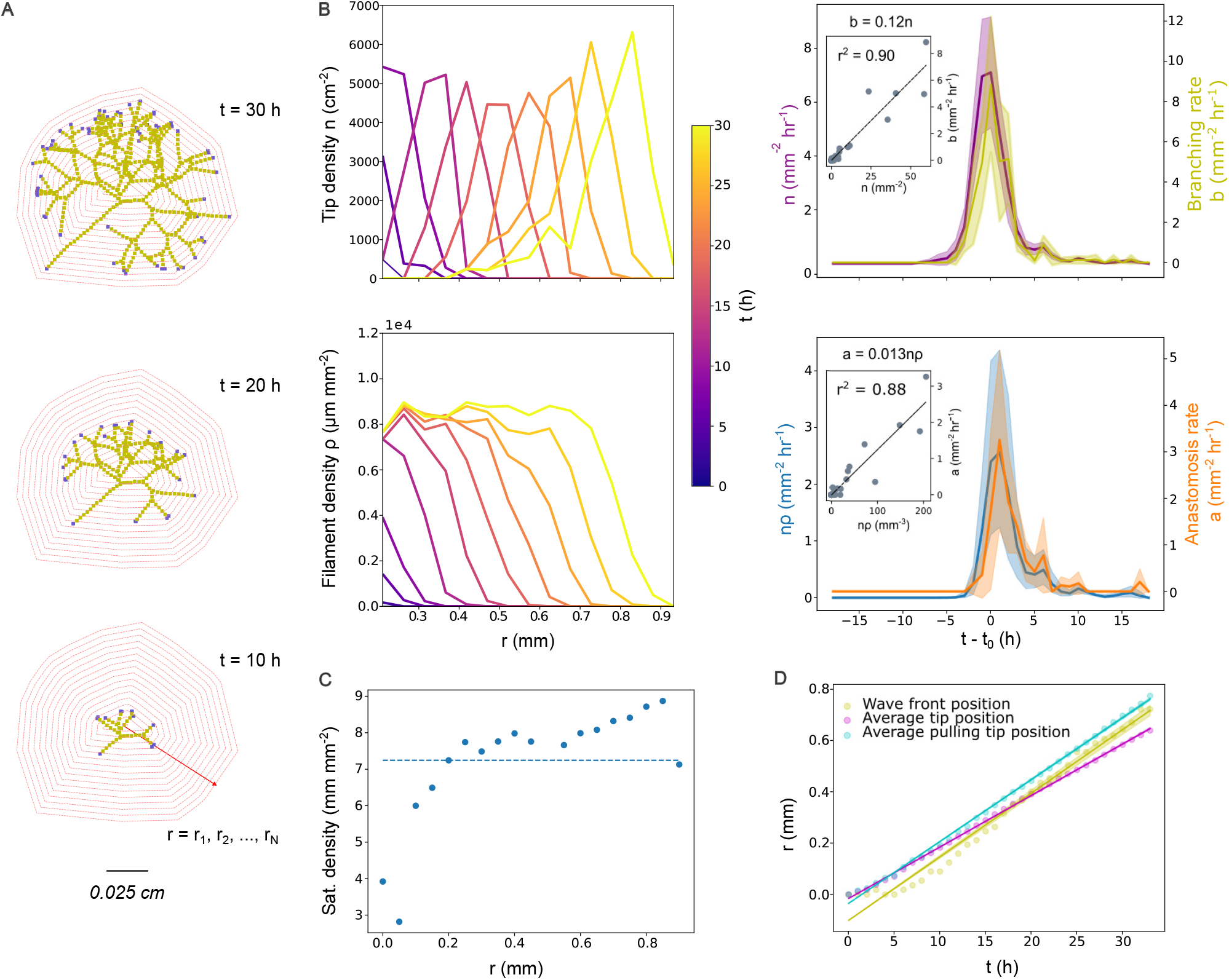
Agent-based simulation of self-regulating traveling wave dynamics. **A** Time series of hyphal network expansion simulated in BMX at 10-hour intervals. Actively growing tips are shown in blue and hyphal filaments in green. Concentric dashed circles indicate radial partitions (*r*_1_, *r*_2_, …, *r*_*N*_ ) used to quantify local segment densities and growth events. **B** Spatial and temporal continuum closures. *Left:* Spatial profiles of active tip density (*n*, top) and hyphal filament density (*ρ*, bottom) over time (purple-to-yellow gradient). *Right:* Temporal dynamics within a representative radial ring. Shaded regions denote standard deviations across stochastic replicates for tip splitting rate *b* (yellow), active tip density *n* (purple), anastomosis rate *a* (orange), and the mass-action product *ρn* (blue). Insets display linear regressions validating the continuum rate closures (*α* = 0.12 h^−1^ and *β* = 13 *µ*m h^−1^). **C** Steady-state filament saturation density (*ρ*_sat_) across successive radial partitions, with the horizontal dashed line denoting the spatial mean. **D** Wave front position over time. The front position (yellow, determined from sigmoid fits to filament density profiles) is compared to the mean position of all tips (pink) and forward-propagating “pulling” tips (cyan). Solid lines indicate linear regressions used to determine wave speeds, with shading representing standard errors. Simulation parameters for the representative run: *l*_max_ = 4.57 *×* 10^−3^ cm, *l*_split_ = 3.89 *×* 10^−3^ cm, *r*_max_ = 1.55 *×* 10^−4^ cm, *p*_*s*_ = 0.15, *p*_*f*_ = 0.01419, *r*_*f*_ = 5.46 *×* 10^−4^ cm, and *W*_*A*_ = 1.11 *×* 10^−4^ cm.

Next, we evaluated whether these macroscopic wave properties are driven by the local growth rules hypothesized in the BARE continuum model (Eq.3). By analyzing the temporal dynamics within a fixed concentric ring as the wave passes through, we found that the rate of apical tip splitting *b* closely tracks the local density of active tips *n*. Similarly, the rate of anastomosis *a* closely tracks the mass-action product *nρ* of tip and filament densities (Fig.1B, right panels). This tight coupling confirms the linear and mass-action rate closures required to link discrete hyphal mechanics to continuum traveling-wave equations.

Across the simulation ensemble, linear regressions confirmed that tip branching scales directly with active tip density (*b* = *αn*, ensemble mean *R*^2^ = 0.95, S1 Fig C). This supports the assumption that new tips emerge near existing growing tips, meaning the branching rate tracks tip density *n* rather than the stationary filament density *ρ*. Conversely, the lower fit for anastomosis rates (*a* = *βnρ*, ensemble mean *R*^2^ = 0.70, S1 Fig C) reflects the breakdown of mass-action assumptions near the expanding front, where filament densities are low and leading tips maintain outward radial alignment from recent branching events. In a representative simulation (*α* = 0.12 h^−1^, *β* = 13 *µ*m h^−1^), branching and fusion rate constants balance in the colony interior in an exact two-to-one ratio (*βρ*_sat_ ≈2*α*≈ 0.24 h^−1^; Fig. 1B insets). This balance stabilizes the interior biomass at a constant saturation density (Fig. 1C), corroborating experimental observations that active tips encounter an integrated average of half the plateau density as the wave passes over them [16].

The simulations confirm that colony expansion on flat agar surfaces operates as a pulled traveling wave (Fig. 1D). The wave speed *c* is governed by the fastest-growing subpopulation of leading tips, which effectively pull the wave forward [18]. These leading tips advance at a velocity faster than the average drift speed (S1 Fig A) and nearly identical to the macroscopic wave speed itself (*c/v*_pull_ = 0.92, S1 Fig B). In contrast, the ensemble mean tip velocity across the entire network is substantially slower than the macroscopic wave front (*c/v*_avg_ = 1.36, S1 Fig B).

Finally, comparing saturation density estimates confirms that the discrete BMX simulations align with the analytical BARE continuum closures (S1 Fig D). Because agent-based simulations are computationally expensive, direct measurements from finite-time BMX runs yield an apparent saturation density (median 1.85 *×* 10^4^ *µ*m mm^−2^) that has not yet reached true steady state. However, the closed-form continuum prediction *ρ*_sat_ = 2*α/β* (median 8.75 *×* 10^4^ *µ*m mm^−2^) and the fitted BARE surrogate value (median 6.2*×* 10^4^ *µ*m mm^−2^) demonstrate where this density ultimately asymptotes. Importantly, all three estimates fall comfortably within the biological range reported for microscopic free-living filamentous fungi (10^4^–10^5^ *µ*m mm^−2^) [14]. Thus, the analytical BARE closures successfully capture the steady-state limits of the BMX dynamics, allowing true saturation densities to be inferred without the need to run computationally prohibitive agent-based simulations to completion.

### Front fluctuations resolve tip transport mechanics

To evaluate how accurately the continuum framework captures discrete hyphal mechanics across growth regimes, we calibrated BARE against three macroscopic quantities of interest extracted from simulated BMX wave profiles: wave amplitude *F*_*n*_ (peak tip density), wave speed *F*_*c*_, and hyphal saturation density *F*_*ρ*_. Jointly minimizing these residuals across simulations yields a three-dimensional Pareto front of rank-1, non-dominated solutions (Fig. 2A). Since the single-objective optima are isolated in distinct extremal regions of this frontier, no single parameter set perfectly minimizes all three residuals. Any global calibration to the BMX dynamics must therefore be an optimal compromise (*F*_opt_) among competing physical constraints.

**Fig 2.**
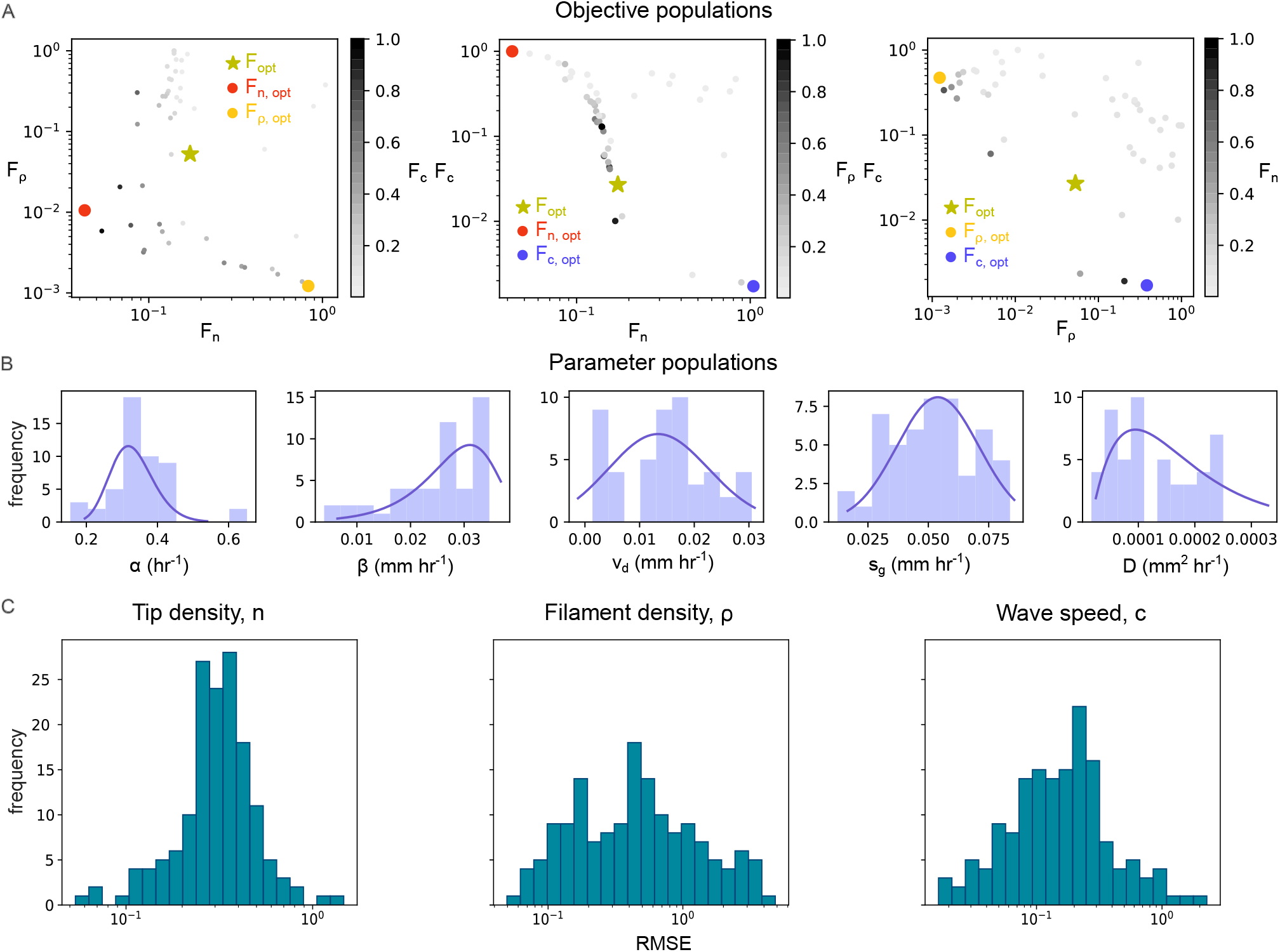
Multi-objective calibration of the continuum surrogate and parameter identifiability. **A** Pair-wise projections of non-dominated (rank-1) solutions forming a three-dimensional Pareto frontier in objective space {*F*_*n*_, *F*_*ρ*_, *F*_*c*_} for the continuum BARE surrogate fit to an example agent-based BMX simulation. Scatter points are colored by the third objective value. The green star indicates the joint compromise optimum (*F*_opt_) minimizing all three weighted residuals, while colored circles mark single-objective optima for wave amplitude *n*_max_ (red), filament saturation density *ρ*_sat_ (yellow), and wave speed *c* (blue). **B** Marginal distributions of rank-1 Pareto solutions in BARE parameter space (*α, β, s*_*g*_, *v*_*d*_, *D*). Solid curves represent fitted distributions used to compute 95% confidence intervals. **C** Distributions of root-mean-square errors (RMSE) for the three objective quantities across the complete ensemble of agent-based model calibrations (N=256).

Pairwise projections of the Pareto front demonstrate that the underlying continuum objectives do not compete equally. The *F*_*n*_ versus *F*_*c*_ projection displays the sharpest and most tightly coupled trade-off (Fig. 2A, middle), forming a narrow, sigmoidal front spanning multiple orders of magnitude from the amplitude optimum (*n*-optimum) to the speed optimum (*c*-optimum). Shading by the third residual (*F*_*ρ*_) indicates that intermediate compromises between amplitude and speed incur higher residuals for the saturation density. The *F*_*n*_ versus *F*_*ρ*_ projection follows an L-shaped frontier where peak tip density trades off against interior filament saturation (Fig. 2A, left). In contrast, the *F*_*ρ*_ versus *F*_*c*_ projection exhibits the weakest trade-off (Fig. 2A, right), appearing as a scattered cloud of rank-1 solutions. This weak coupling confirms that wave speed and interior filament density operate through largely distinct macroscopic constraints, though fitting both simultaneously remains constrained by amplitude fidelity.

Mapping the Pareto solutions back into BARE model space reveals a structurally anisotropic parameter landscape (Fig. 2B, S2 Fig). The system is stiff along combinations that dictate net kinetic rates: the marginal distributions for the branching rate *α*, anastomosis rate *β*, and elongation velocity *s*_*g*_ are unimodal and tightly localized. These yield narrow 95% confidence intervals, reflecting robust identifiability of the theoretical saturation density 2*α/β*. Conversely, directed drift velocity *v*_*d*_ and effective tip diffusivity *D* align along an unconstrained, sloppy manifold [47]. Since the deterministic wave front propagates at the pulled-front speed 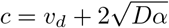, any pair (*v*_*d*_, *D*) satisfying this relationship produces an identical wave speed for a given branching rate *α*. This continuous degeneracy constitutes the mechanistic origin of the loose *F*_*ρ*_–*F*_*c*_ Pareto trade-off, demonstrating that mean-field wave profiles alone cannot disentangle directed tip elongation from dispersive hyphal exploration.

Resolving this structural non-identifiability requires information beyond the deterministic wave profile. While finite-tip demographic noise induces a stochastic velocity correction Δ*c*, this shift represents only a 1–3% reduction unless the effective number of tips at the leading edge is very small (*N*_*e*_ ∼ *O* (1)), making it difficult to distinguish from stochastic variance in measured speeds across BMX simulations. We instead resolved the (*v*_*d*_, *D*) manifold by measuring the front-wandering diffusivity *D*_*f*_ from the growth in mean squared displacement of the front position across stochastic realizations (Eq. 21, S3 Fig A). Applying the theoretical front-wandering scaling relation (Eq. 20) to empirical *D*_*f*_ measurements uniquely determines the effective diffusivity *D*, after which the directed drift velocity *v*_*d*_ follows directly from the measured wave speed (Eq. 14). Across all simulations, the value of *D* isolated by front wandering consistently accumulated near the lower bound of the confidence interval estimated from the deterministic PDE (S3 Fig B). This systematic downward shift reflects a fundamental biological distinction between leading-edge and interior dynamics. While the mean-field PDE assumes uniform diffusion throughout the mycelium, leading tips at the expanding front exhibit strong directional persistence and negative autotropism—a self-avoidance mechanism that optimizes the spatial occupation of substrate [37]—resulting in a lower effective diffusivity at the front than in the isotropic interior.

Overall, front speed and peak tip density were the most accurately recovered quantities, yielding a mean root-mean-square error (RMSE) of 0.24 for *F*_*c*_ and 0.34 for *F*_*n*_ (Fig. 2C). In comparison, saturation density showed greater residual spread (mean RMSE 0.88). This difference arises from two compounding factors: front tracking provides a less noisy observable than local filament density estimates (S4 Fig), and the geometry of the Pareto front allows saturation density to vary considerably while speed and amplitude remain tightly optimized near their minima (Fig. 2A, middle). In summary, while macroscopic measurements (wave speed, peak tip density, and saturation density) constrain the overall shape and speed of the expanding colony, observing stochastic front fluctuations is necessary to separate directed hyphal elongation (*v*_*d*_) from dispersion (*D*).

### Surrogate models bridge discrete and continuum scales

To connect discrete hyphal growth rules to macroscopic wave dynamics across parameter space, we constructed a continuous mapping from the seven microscopic BMX parameters (Table 2) to the five continuum BARE parameters (Table 1). For any input combination of BMX parameters, the surrogate defines a confidence region of consistent continuum BARE model parameters (Fig. 3A, B).

**Fig 3.**
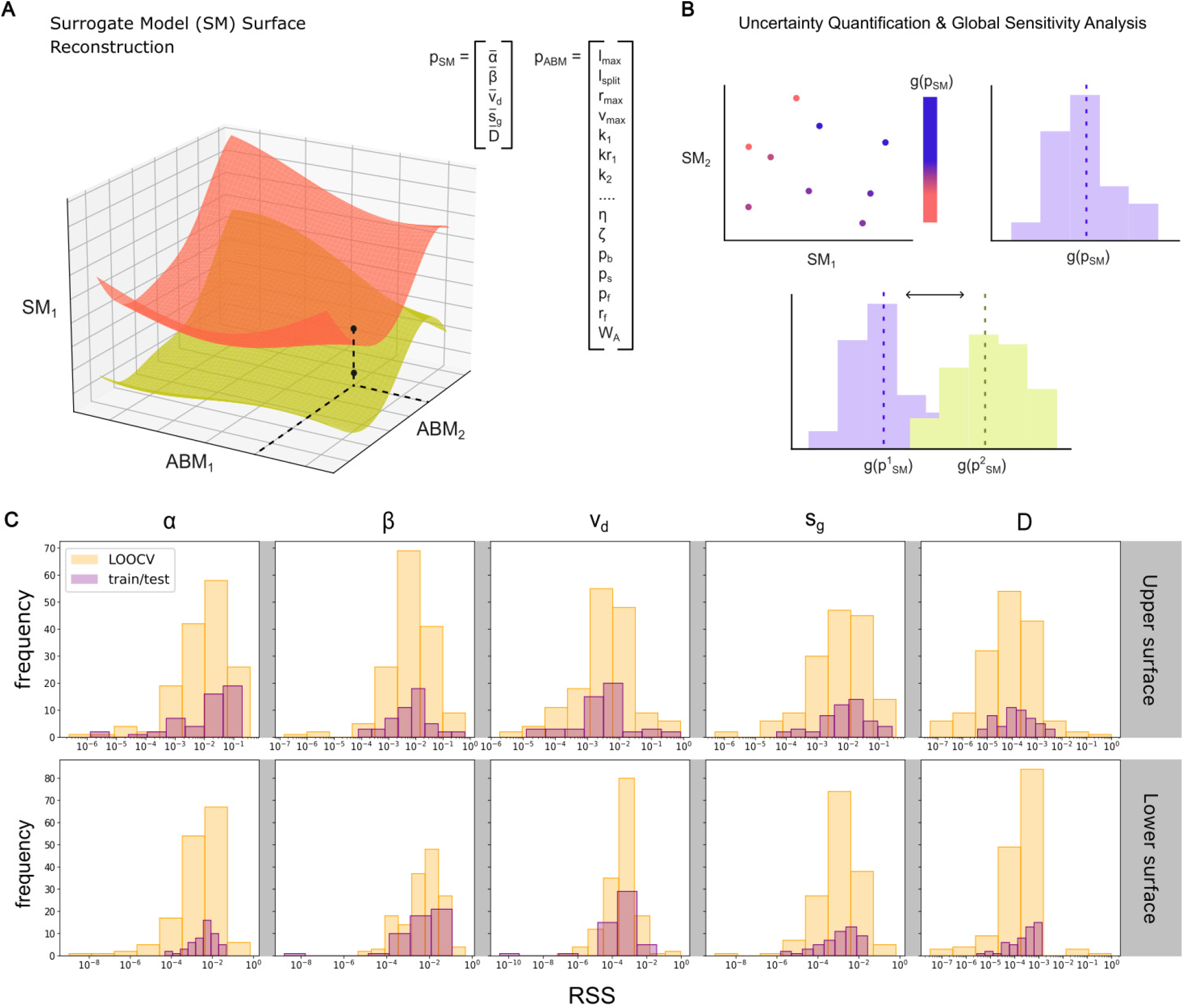
Continuum surrogate construction and cross-validation. **A** Continuous mapping from agent-based (BMX) input parameters (*p*_ABM_) to continuum (BARE) surrogate parameters (*p*_SM_). For each continuum model parameter, radial basis functions (RBFs) interpolate the upper (orange) and lower (yellow) 95% confidence bounds across the agent-based parameter domain. For visualization, two agent-based parameters and one continuum parameter are shown; across the full model, each bounding hypersurface is a seven-dimensional function over the varied BMX inputs. **B** Continuum parameters are sampled uniformly within the bounded hyperrectangle defined by the upper and lower surfaces (top left), generating a distribution for the macroscopic output *g*(*p*_SM_) whose mean represents the marginalized surrogate output (top right). Evaluating shifts across output distributions under parameter variations enables computationally efficient global sensitivity analysis of the agent-based model (bottom). **C** Residual sum of squares (RSS) error distributions for upper and lower bounding surfaces across all five continuum BARE parameters (*α, β, v*_*d*_, *s*_*g*_, *D*). Histograms compare leave-one-out cross-validation on the training set (70%, yellow) with predictions on unseen inputs from a 30% held-out test set (purple).

This mapping generalizes accurately across the sampled domain, with squared residuals for both leave-one-out cross-validation (LOOCV) on the training set and a 30% held-out test set concentrated between 10^−3^ and 10^−2^ (Fig. 3C). The close overlap of the training and test error distributions indicates that the surrogate generalizes to unseen BMX inputs without substantial loss of accuracy, with at most a modest residual increase for specific parameters such as the branching rate *α*. By marginalizing over this parameter mapping uncertainty, the continuum surrogate allows the system’s behavior and its underlying ‘stiff and sloppy’ directions to be evaluated continuously [47]. This establishes an analytically tractable, computationally efficient foundation for global sensitivity analysis across the full space of hyphal growth behaviors.

### Distinct hyphal parameters govern wave speed and saturation density

Global sensitivity analysis reveals that macroscopic colony dynamics are driven primarily by four of the seven varied microscopic parameters in the agent-based model: maximum segment length (*l*_max_), hyphal radius (*r*_max_), tip-splitting probability (*p*_*s*_), and fusion radius (*r*_*f*_ ) (Fig. 4). Both variance-based (Sobol) and distribution-based (PAWN) sensitivity methods yield identical parameter importance rankings across all continuum outputs. Importantly, these microscopic parameters partition into two largely separate groups that independently regulate filament saturation density (*ρ*_sat_) and wave speed (*c*).

**Fig 4.**
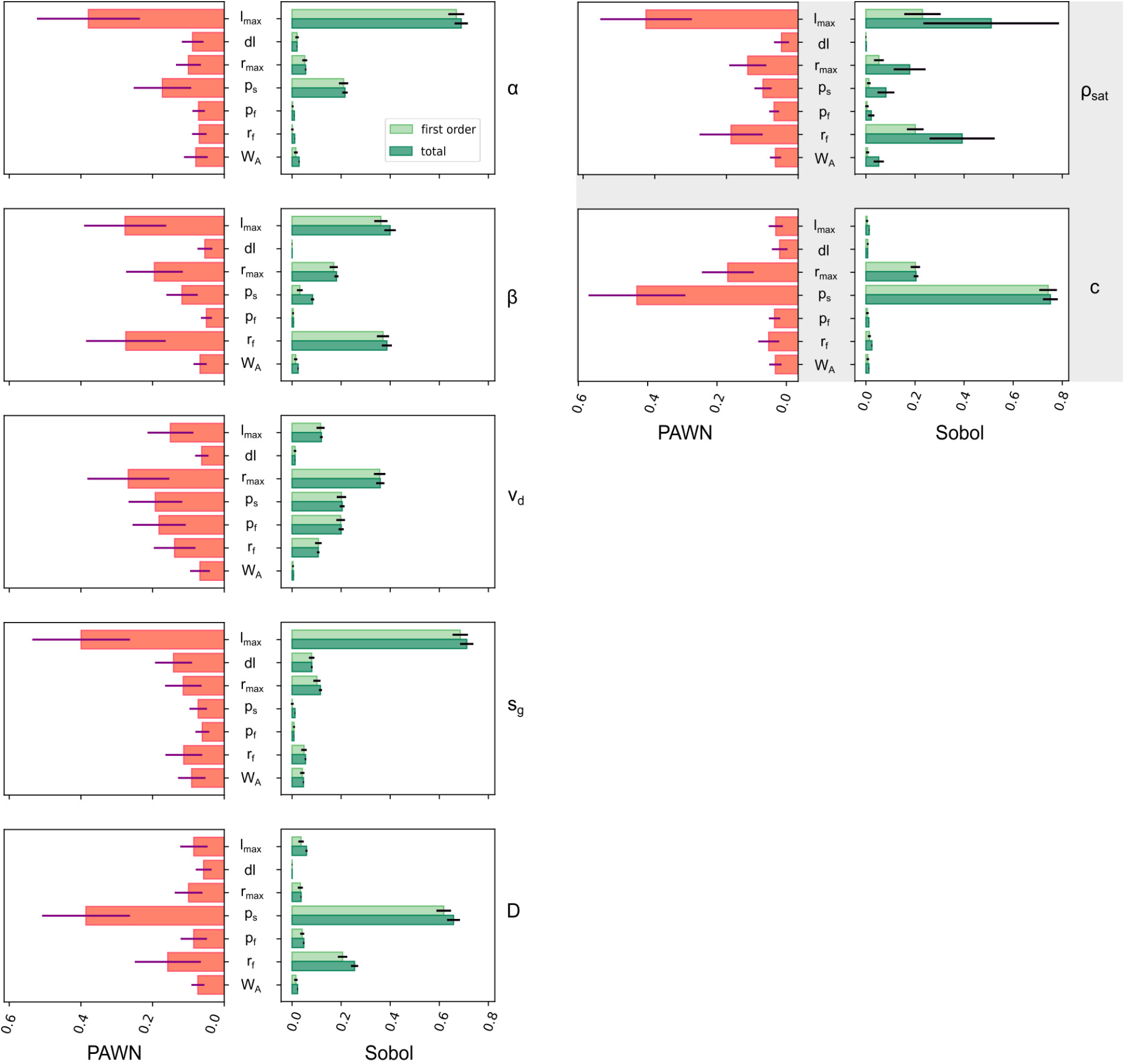
Global sensitivity analysis of continuum traveling wave observables to hyphal parameters. Comparison of distribution-based (PAWN, left column) and variance-based (Sobol, right column) global sensitivity indices evaluating the influence of seven agent-based (BMX) model parameters (*l*_max_, *dl, r*_max_, *p*_*s*_, *p*_*f*_, *r*_*f*_, *W*_*A*_) across continuum (BARE) parameters (*α, β, v*_*d*_, *s*_*g*_, *D*) and emergent macroscopic traveling-wave observables (*ρ*_sat_, *c*; shaded panels). For PAWN indices, bars represent the median Kolmogorov–Smirnov statistic, and error bars denote standard deviations across bootstrap resamples. For Sobol indices, light and dark green bars represent first-order (*S*_*i*_) and total-order (*S*_*T i*_) sensitivity indices, respectively, with error bars indicating 95% bootstrap confidence intervals.

Maximum segment length *l*_max_ is the dominant driver of saturation density *ρ*_sat_ (accounting for ∼ 50% of total variance; Fig. 4, top right). Because hyphae evaluate apical splitting and fusion interactions upon completing each growth segment, a shorter *l*_max_ increases the spatial frequency of potential splitting events per unit length, generating a denser network within the colonized interior. The fusion radius (*r*_*f*_ ) and hyphal radius (*r*_max_) exert the next strongest influences on *ρ*_sat_ (total Sobol indices ∼ 0.40 and ∼ 0.20, respectively). Because anastomosis occurs when searching tips encounter existing filaments, *r*_*f*_ and *r*_max_ act as geometric thresholds that expand the physical contact regions facilitating fusion. Furthermore, the large difference between first- and total-order Sobol indices for *ρ*_sat_ reveals strong interaction effects among *l*_max_, *r*_*f*_, and *r*_max_, indicating that interior packing is a collective property of network geometry rather than a simple sum of isolated hyphal traits.

In contrast, the wave speed *c* is predominantly governed by the probability of tip-splitting *p*_*s*_, which accounts for approximately 75% of the total variance, alongside a secondary contribution from hyphal radius *r*_max_ ( ∼ 20%; Fig. 4, bottom right). Under the pulled-front dynamics of the expanding margin, colony expansion speed depends critically on the rate and spatial orientation of leading tips. Higher splitting rates increase the number of active tips at the periphery, raising the statistical likelihood that a subset of leading tips aligns directly along the outward growth vector to pull the front forward. Notably, a short segment length (*l*_max_) does not inherently slow wave propagation: when tip-splitting and fusion probabilities are low, hyphal tips simply continue to elongate forward across successive segment intervals without splitting, leaving wave speed unaffected.

This distinction between interior network densification and directed front motion is reflected in the individual continuum transport parameters. While the branching rate *α* and mean elongation rate *s*_*g*_ are both dictated by segment length *l*_max_ (*>* 70% of variance), directed drift velocity *v*_*d*_ shows distributed sensitivity across almost all mechanical parameters: hyphal radius (*r*_max_), splitting probability (*p*_*s*_), fusion probability (*p*_*f*_ ), segment length (*l*_max_), and fusion radius (*r*_*f*_ ) (Fig. 4, left). Thus, rather than scaling with any single microscopic trait, drift velocity captures how local, isotropic growth rules collectively break symmetry to generate globally coordinated outward expansion.

This mechanistic separation is further reinforced by the behavior of the effective diffusivity *D*. While the fusion radius *r*_*f*_ accounts for ∼ 30% of the variance in *D*, this variance does not propagate to the wave speed *c* (Fig. 4). Diffusivity across the parameter space is highest when *r*_*f*_ is small, as hyphae must wander further across space to encounter fusion partners without being annihilated upon colliding with trailing filaments. This decoupling reinforces the necessity of using leading-edge front wandering (*D*_*f*_ ) to constrain transport: wave speed *c* is governed strictly by the leading tips at the uncolonized front where fusion collisions are virtually absent, whereas interior diffusion is strongly shaped by anastomosis.

Finally, variations in the post-split segment length difference (*dl*) and the width of the hyphal attraction region (*W*_*A*_) show negligible sensitivity across all macroscopic continuum properties (Fig. 4). Because *W*_*A*_ acts only over a narrow region near the hyphal surface, its force is easily overcome by tip elongation. At the sparse leading edge, tips rarely come close enough to feel this attraction, while in the interior, fusion is governed primarily by the larger geometric radius *r*_*f*_ . Thus, while negative autotropism maintains spacing at the expanding front, localized attractive tropisms (*W*_*A*_) and differences in post-split segment lengths (*dl*) are rendered negligible by the primary rules of hyphal elongation, apical splitting, and fusion.

### Hyphal trait combinations map onto explorer–exploiter growth strategies

To evaluate how specific combinations of hyphal traits structure the balance between interior biomass density and outward expansion speed, we mapped the ratio of normalized filament saturation density to wave speed 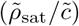 across all pairwise parameter projections (Fig. 5B). High values of this ratio identify the exploiter strategy—characterized by dense, slower-moving networks—while low values identify the fast, sparse explorer strategy [14].

**Fig 5.**
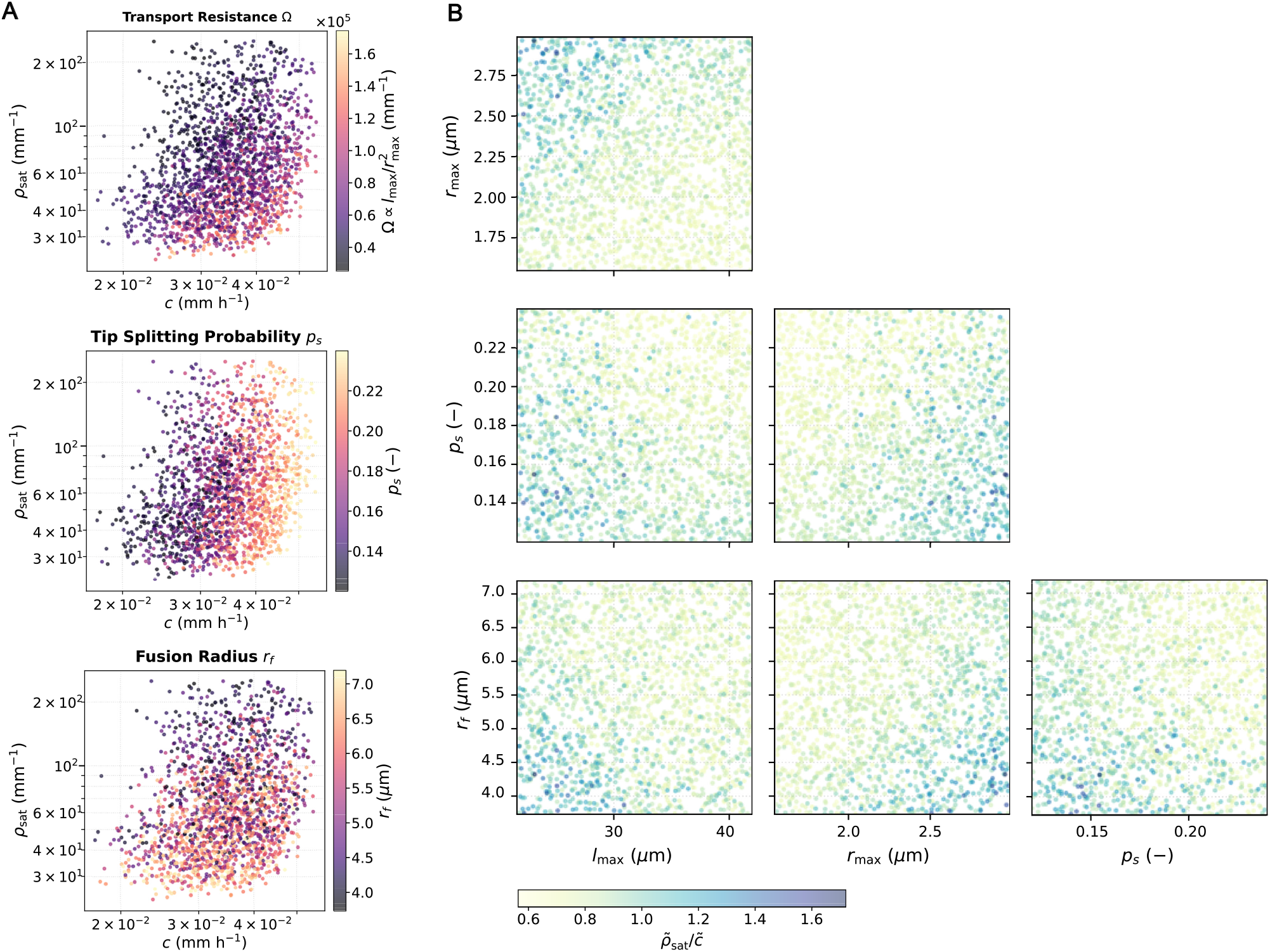
Fungal growth strategies across continuum and agent-based parameter space. **A** Relationship between wave speed (*c*) and saturation density (*ρ*_sat_), colored by transport resistance 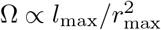, top), tip-splitting probability (*p*_*s*_, middle), and fusion radius (*r*_*f*_, bottom). **B** The mechanically permissible (potential) trait space generated across variations in hyphal parameters: segment length (*l*_max_), hyphal radius (*r*_max_), tip-splitting probability (*p*_*s*_), and fusion radius (*r*_*f*_ ). Points represent individual agent-based simulations colored by the normalized ratio 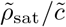 (with wave speed and saturation density each normalized to the range 1–2). This ratio captures a morphological spectrum spanning from fast, sparse exploration (pale yellow, 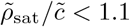 to slow, dense exploitation (deep blue, 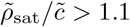.

Notably, the continuum and agent-based frameworks permit a broad potential trait space, generating networks that can theoretically achieve both high expansion speeds and high saturation densities simultaneously (Fig.5A, top right quadrant of scatter plots). However, when isolating the distinct ecological phenotypes, the pairwise projections demonstrate that they occupy localized, opposing geometric regions within this broader parameter space (Fig.5B).

The dense exploiter regime 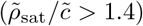 is realized when short hyphal segments (*l*_max_) and small fusion radii (*r*_*f*_ ) coincide with thick hyphae (*r*_max_) and lower apical splitting probabilities (*p*_*s*_). This structural configuration yields networks with low geometric transport resistance 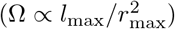. This indicates that dense exploiter morphologies are inherently characterized by low internal resistance, whereas fast explorer networks are geometrically unconstrained by it (Fig. 5A, top).

Conversely, the explorer regime emerges when long hyphal segments and high apical splitting rates prioritize the generation and forward propagation of leading tips rather than local mesh consolidation. By tuning these four primary hyphal growth parameters, the discrete biophysical framework successfully spans the continuous morphological spectrum between exploration and exploitation observed across fungal foraging strategies [40, 48].

## Discussion

Bridging microscopic hyphal growth rules to macroscopic colony expansion helps resolve how fungal networks self-organize across physical scales [1–3]. Our multiscale framework demonstrates that mycelial colony expansion operates as a pulled traveling wave [18] governed by distinct physical processes at the leading edge versus the dense network interior. Methodologically, the stochastic BARE continuum surrogate provides a computationally efficient bridge between these discrete microscopic simulations and macroscopic reaction-diffusion models [31, 49]. By measuring demographic fluctuations at the advancing front (*D*_*f*_ ) to isolate effective diffusion (*D*) from directed drift (*v*_*d*_), this framework resolves a structural non-identifiability in the deterministic wave speed using independent physical observables [22, 32].

From a biophysical perspective, the model shows how local hyphal growth rules dictate transport constraints and shape the 3D colony profile. At the advancing front, wave speed is driven primarily by apical tip splitting. More frequent splitting increases the statistical likelihood that a subset of active leading tips aligns directly along the outward radial growth direction to pull the front forward. This positive relationship between splitting frequency and expansion speed agrees with empirical studies showing that hyphal reach scales directly with splitting intensity [50]. Furthermore, hyphal tips diffuse more slowly at the expanding margin than in the interior. This mathematical decoupling matches microscopic observations of self-avoidance (negative autotropism) in 2D, where exploring tips mechanically repel each other to evenly occupy uncolonized substrate [37].

Behind this leading edge, the colony interior naturally stabilizes when apical tip splitting balances fusion in an exact two-to-one ratio (*βρ*_sat_ = 2*α*). This ratio emerges because the active tip population is concentrated at the advancing front, where it encounters an average of half the final saturation density (*ρ*_sat_*/*2) as the wave passes. This verifies that mycelial networks self-regulate biomass density through local tip–filament collisions. It also confirms that density saturation under replete growth conditions is not driven by environmental carrying capacity, but represents a specific growth strategy under fungal control. Indeed, recent empirical studies of traveling-wave fungal growth demonstrate that this saturation density remains remarkably invariant despite changes in carbon availability, or even when advancing wavefronts collide [16]. Because the theoretical saturation density contains neither expansion speed nor diffusion, interior biomass remains strictly independent of front velocity. This allows the fungus to dictate its own spatial densification through anastomosis regardless of how fast it explores new territory. Additionally, separating front expansion from interior network formation suggests that the continuum framework could be expanded by incorporating a density-dependent diffusion rate, *D*(*ρ*), explicitly capturing how hyphal tip movement slows down as the network interior becomes crowded [51].

By isolating pure hyphal mechanics, this framework highlights a fundamental separation between potential and realized trait spaces. Our results demonstrate that the empirical explorer–exploiter trade-off—the choice between rapid spatial exploration and local densification [16]—is not an unavoidable mechanical constraint. Because wave velocity is primarily governed by apical tip splitting while saturation density is dictated by segment length, pure geometry permits a large potential trait space, including theoretical colonies that are both fast and dense. Nevertheless, our mechanical baseline grounds specific geometric trait correlations observed in living fungi. In the trait ordination of microscopic fungi [14], the dominant axis opposes dense networks with high edge length density and meshedness against exploratory ones with high edge length, edge width, and root-to-tip transport efficiency. Our simulations capture these morphological trade-offs. First, confirming the empirical negative correlation between individual edge dimensions and overall network density, we find that both shorter segment lengths and narrower hyphal radii drive higher saturation densities. Furthermore, transport resistance 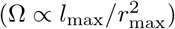 provides a physical mechanism for why edge length and width strongly covary in exploratory networks. When fungi elongate their segments to explore space, they must simultaneously thicken their hyphae; because radius is squared in the denominator, this widening offsets the resistance penalty of long edges to maintain high transport efficiency. Yet, because pure mechanics theoretically allow segment length, radius, and tip splitting to combine into fast-and-dense networks, the restricted explorer–exploiter spectrum observed in soils must arise from external limits. Building dense biomass while fueling rapid expansion carries carbon construction costs. Thus, while local growth rules define a broad baseline of physically possible forms, finite metabolic budgets—coupled with host-dictated nutrient exchange [17] and defense against grazing [40, 48]—ultimately restrict fungi to the narrow realized spectrum of foraging strategies observed in nature.

Ultimately, these findings offer a practical way to improve how large-scale biogeochemical models represent fungal colony growth. Instead of mechanistically capturing traveling wave dynamics, current spatially-explicit grid-based models simulate colonization by applying a generic solid-phase diffusion coefficient to a bulk biomass pool [52]. Similarly, they rely on arbitrarily tuned carrying capacities (*K*) to cap local biomass accumulation [53]. While agent-based models easily capture the explicit hyphal traits that actually drive these processes, their immense computational cost prevents their use at ecosystem scales. The BARE continuum model resolves this bottleneck by translating discrete hyphal traits into mathematically simple macroscopic parameters. Specifically, the traveling wave speed (*c*) provides a colonization velocity grounded directly in apical elongation and branching rates; by applying this speed as a trait-driven advective flux, models can explicitly simulate both sub-grid patch expansion and grid-to-grid colonization, replacing the generic solid-phase diffusion currently used in the advection-dispersion transport schemes of next-generation soil biogeochemistry models [54]. Simultaneously, our algebraically derived saturation density (*ρ*_sat_) transforms how models handle carrying capacity. By multiplying *ρ*_sat_ by hyphal cross-sectional area and tissue carbon density, biogeochemical reaction networks can calculate the maximum biomass carbon a specific soil volume can physically hold [55]. Consequently, carrying capacity ceases to be an arbitrarily tuned model parameter and instead emerges as a strictly bounded intrinsic fungal growth strategy defined by phylogenetically conserved morphological traits [14, 56]. By providing these explicit physical limits as computationally cheap parameterizations, this framework establishes an operational foundation to translate soil taxonomic data and microscopic fungal architecture into the macroscopic colonization rates that drive terrestrial carbon cycling [57].

## Acknowledgments

This work was supported by the Laboratory Directed Research and Development Program of Lawrence Berkeley National Laboratory under U.S. Department of Energy Contract No. DE-AC02-05CH11231, and by the U.S. Department of Energy, Office of Science, Office of Workforce Development for Teachers and Scientists (WDTS) under the Science Undergraduate Laboratory Internship (SULI) program. The authors thank Andrew T. Myers (Computing Sciences Area, Lawrence Berkeley National Laboratory, Berkeley, CA) and Bruce J. Palmer (Physical and Computational Sciences Directorate, Pacific Northwest National Laboratory, Richland, WA) for their technical guidance and helpful discussions regarding the Biological Modeling and interface eXchange (BMX) open-source software package (BSD-2-Clause license).

## Author contributions

Conceptualization: Gianna L. Marschmann. Methodology: Gianna L. Marschmann, Lauren Yan. Software: Lauren Yan. Formal analysis: Lauren Yan. Investigation: Lauren Yan. Funding acquisition: Gianna L. Marschmann. Project administration: Gianna L. Marschmann. Supervision: Gianna L. Marschmann. Visualization: Lauren Yan, Gianna L. Marschmann. Writing – original draft: Lauren Yan, Gianna L. Marschmann. Writing – review & editing: Lauren Yan, Gianna L. Marschmann.

## Code availability

All source code, BMX configuration files, and analysis scripts required to reproduce the results presented in this manuscript are freely available in a public GitHub repository at *https://github.com/giannamars/bmx-fungal-continuum-surrogates*. The accompanying simulation datasets are available from the corresponding author upon reasonable request.

## Supporting information

**S1 Fig.**
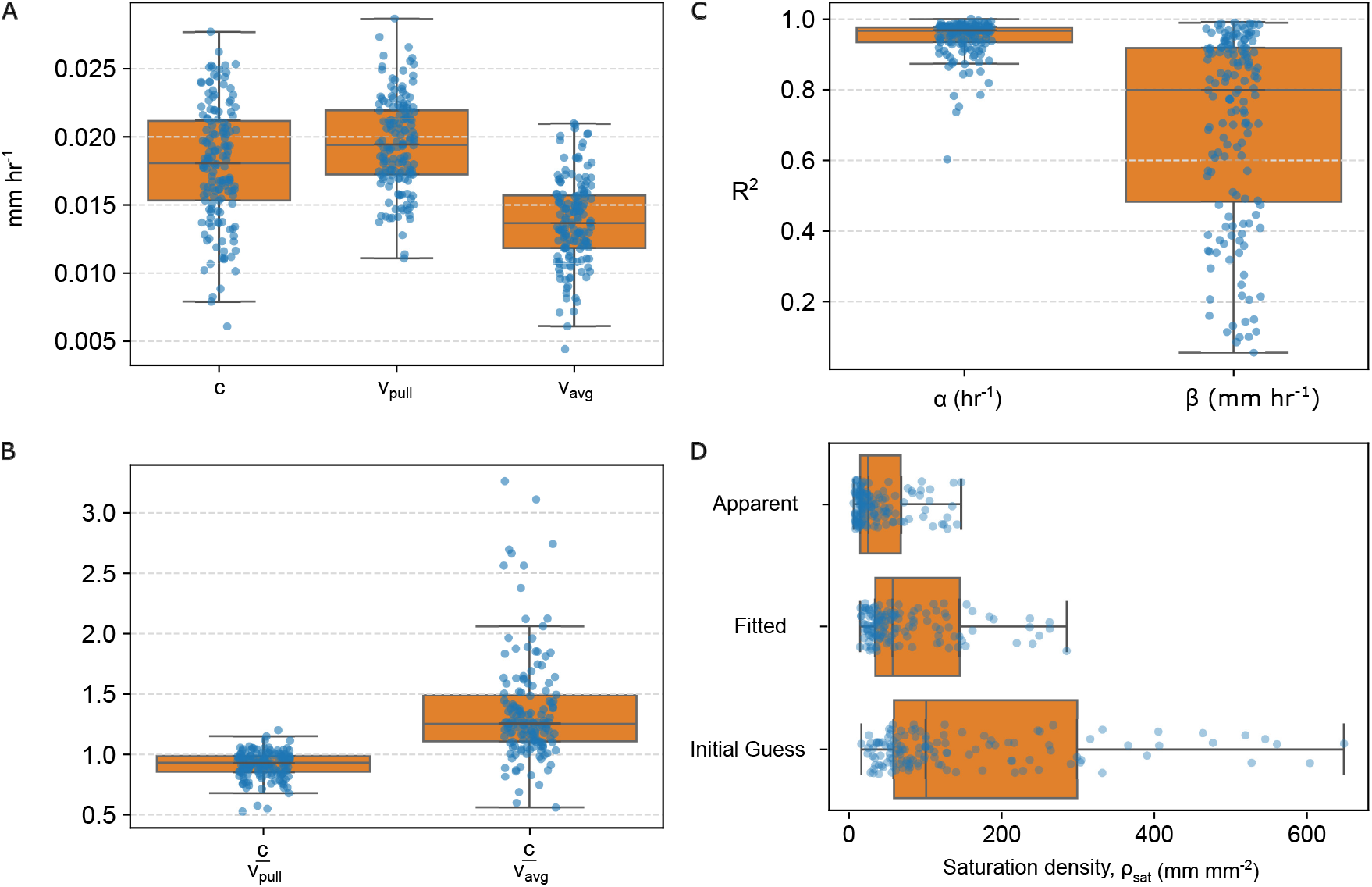
Validation of continuum closures and pulled wave dynamics. **A** Macroscopic wave speed (*c*), average growth velocity of leading tips (*v*_pull_), and ensemble average tip velocity (*v*_avg_) across BMX simulations. **B** Ratios between the macroscopic wave speed and the two tip velocities. **C** Goodness-of-fit (*R*^2^) for the linear regressions of apical tip splitting (*b* = *αn*) and anastomosis (*a* = *βnρ*) rates across the simulation ensemble. **D** Comparison of saturation density *ρ*_sat_ estimates across BMX simulations: apparent density measured directly from finite-time BMX runs, fitted density from the BARE surrogate, and the analytical closed-form prediction (*ρ*_sat_ = 2*α/β*). Across **A**–**D**, the top and bottom of each boxplot mark the 75th and 25th percentiles, the horizontal line inside marks the median, and the whiskers span the data range excluding outliers.

**S2 Fig.**
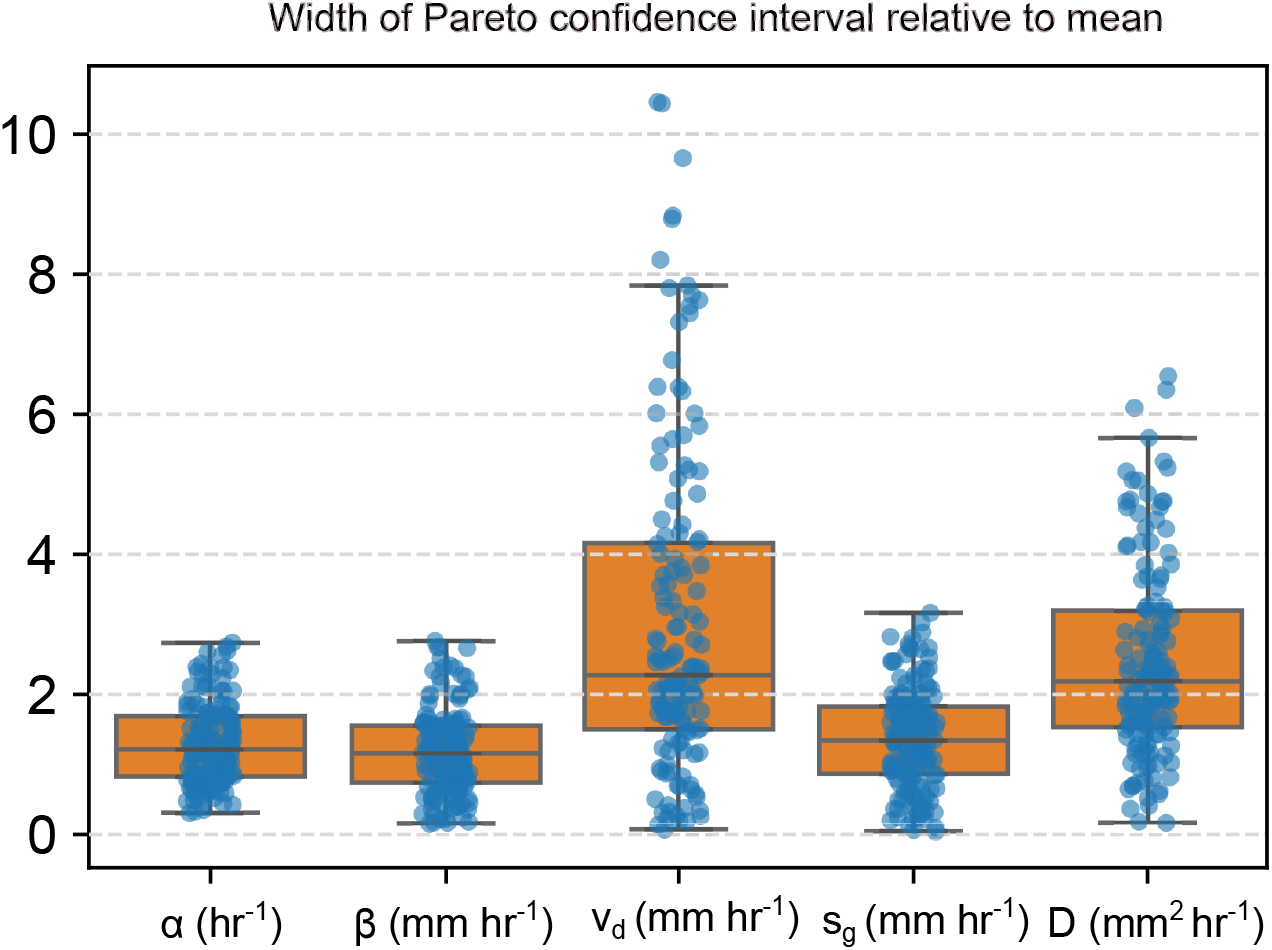
Identifiability of continuum parameters across the Pareto front. Each boxplot displays the distribution of 95% confidence interval widths—normalized by the mean of the Pareto-optimal population—for each continuum parameter (*α, β, v*_*d*_, *s*_*g*_, *D*) across the BMX simulation ensemble. The top and bottom of each boxplot mark the 75th and 25th percentiles, the horizontal line inside marks the median, and the whiskers span the data range including outliers.

**S3 Fig.**
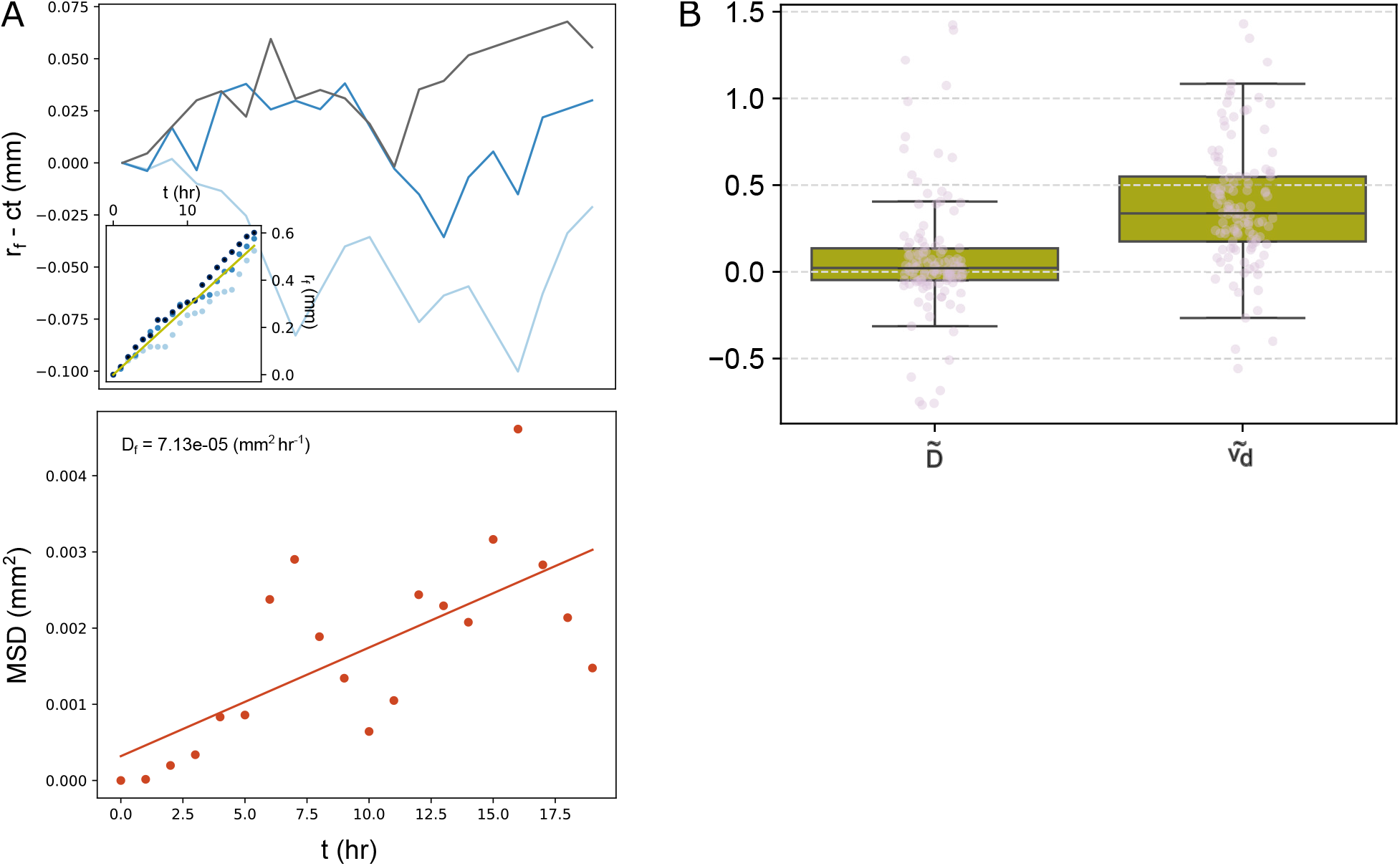
Resolving structural non-identifiability via stochastic front wandering. **A** Estimation of front-wandering diffusivity (*D*_*f*_ ) for a representative BMX simulation. The top inset displays the absolute position of the wave front over time across three stochastic replicates, with the solid green line indicating the deterministic advance at the mean wave speed. The main upper plot shows each replicate’s spatial deviation from this mean over time. The lower scatter plot tracks the growth of the mean squared displacement (MSD) of the front across the replicates, with the solid line representing the linear regression used to infer *D*_*f*_ . **B** Distributions of the effective tip diffusivity (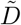) and directed drift velocity (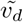) resolved via front wandering across the simulation ensemble. Values for each simulation are normalized relative to their 95% confidence intervals derived from the deterministic Pareto front, where 0 represents the lower bound and 1 represents the upper bound. The top and bottom of each boxplot mark the 75th and 25th percentiles, the horizontal line inside marks the median, and the whiskers span the data range including outliers.

**S4 Fig.**
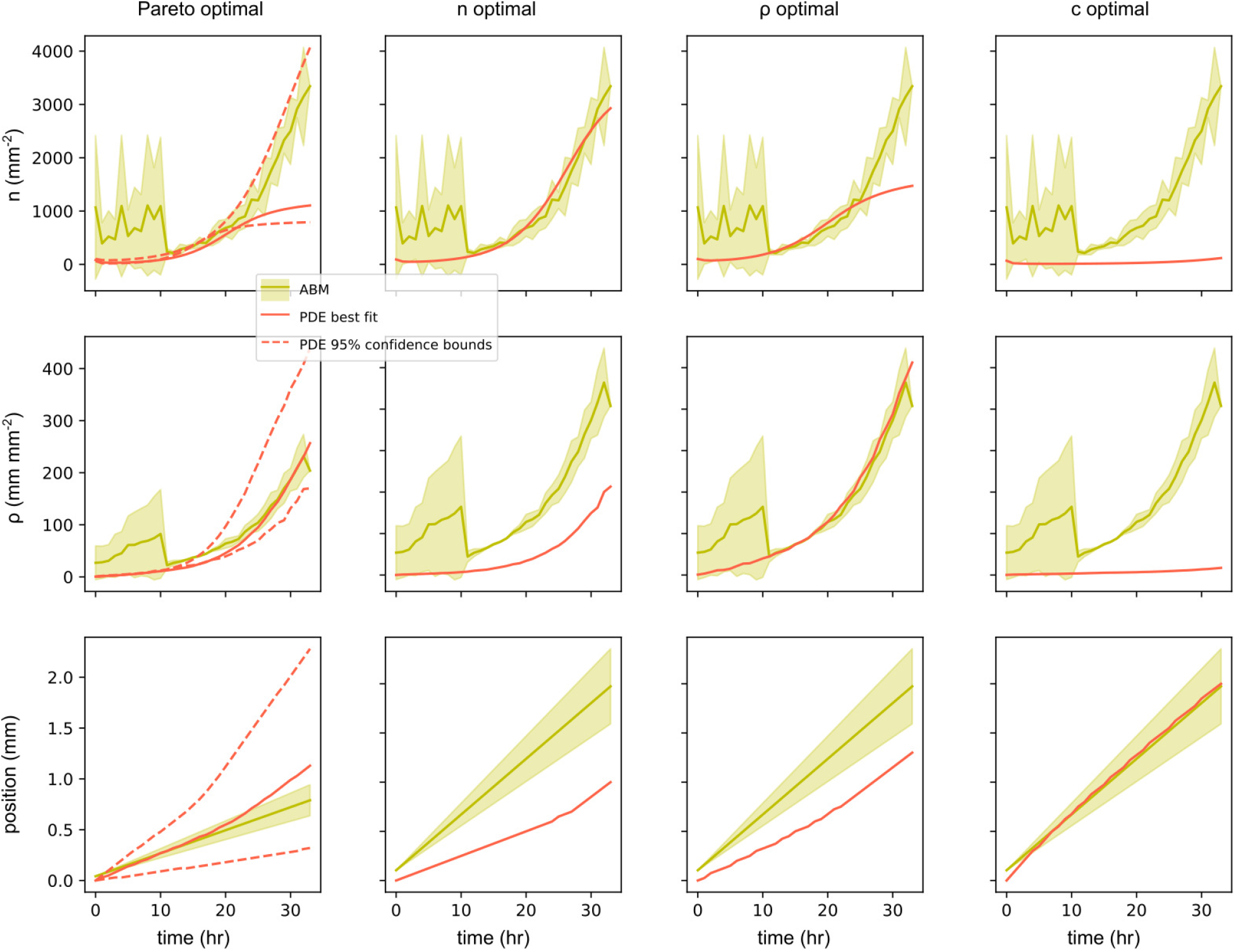
Comparison of the Pareto-optimal compromise to single-objective optima. Each column shows the BARE continuum model fit to the macroscopic BMX simulation data when optimized jointly for all three objectives (the Pareto compromise) versus optimized individually for peak tip density (*n*), saturation density (*ρ*), or wave speed (*c*). BARE continuum predictions are shown in red (solid line for the best fit; dashed lines for the 95% confidence interval). Macroscopic BMX data are plotted in green (solid line for the median across three stochastic replicates; shaded region for the 95% confidence interval). The Pareto-optimal solution represents the best global compromise, identifying a parameter vector where no single objective can be further improved without degrading the fit of the others.

## References

1. Camazine S, Deneubourg JL, Franks NR, Sneyd J, Theraula G, Bonabeau E. Self-organization in biological systems. Princeton, New Jersey: Princeton University Press; 2001. doi:10.2307/j.ctvzxx9tx.

2. Weitz JS, Benfey PN, Wingreen NS. Evolution, interactions, and biological networks. PLoS biology. 2007;5(1):e11.

3. Hallatschek O, Datta SS, Drescher K, Dunkel J, Elgeti J, Waclaw B, et al. Proliferating active matter. Nature Reviews Physics. 2023;5(7):407–19.

4. Money NP. Insights on the mechanics of hyphal growth. Fungal Biology Reviews. 2008;22(2):71–6.

5. Daskalov A, Heller J, Herzog S, Fleißner A, Glass NL. Molecular mechanisms regulating cell fusion and heterokaryon formation in filamentous fungi. Microbiology spectrum. 2017;5(2):10–1128.

6. Roper M, Seminara A. Mycofluidics: the fluid mechanics of fungal adaptation. Annual Review of Fluid Mechanics. 2019;51(1):511–38.

7. Harris SD. Branching of fungal hyphae: regulation, mechanisms and comparison with other branching systems. Mycologia. 2008;100(6):823–32.

8. Fricker MD, Heaton LL, Jones NS, Boddy L. The mycelium as a network. The fungal kingdom. 2017:335–67.

9. Steinberg G. Hyphal growth: a tale of motors, lipids, and the Spitzenkorper. Eukaryotic cell. 2007;6(3):351–60.

10. Riquelme M, Yarden O, Bartnicki-Garcia S, Bowman B, Castro-Longoria E, Free SJ, et al. Architecture and development of the Neurospora crassa hypha–a model cell for polarized growth. Fungal biology. 2011;115(6):446–74.

11. Schmideder S, Müller H, Barthel L, Friedrich T, Niessen L, Meyer V, et al. Universal law for diffusive mass transport through mycelial networks. Biotechnology and Bioengineering. 2021;118(2):930–43.

12. Bebber DP, Hynes J, Darrah PR, Boddy L, Fricker MD. Biological solutions to transport network design. Proceedings of the Royal Society B: Biological Sciences. 2007;274(1623):2307.

13. Davidson FA. Mathematical modelling of mycelia: a question of scale. Fungal Biology Reviews. 2007;21(1):30–41.

14. Aguilar-Trigueros CA, Boddy L, Rillig MC, Fricker MD. Network traits predict ecological strategies in fungi. ISME communications. 2022;2(1):2.

15. Dikec J, Olivier A, Bobée C, D’angelo Y, Catellier R, David P, et al. Hyphal network whole field imaging allows for accurate estimation of anastomosis rates and branching dynamics of the filamentous fungus Podospora anserina. Scientific reports. 2020;10(1):3131.

16. Oyarte Galvez L, Bisot C, Bourrianne P, Cargill R, Klein M, van Son M, et al. A travelling-wave strategy for plant–fungal trade. Nature. 2025;639(8053):172–80.

17. Bisot C, Galvez LO, Kahane F, Van Son M, Turcu B, Broekman R, et al. Carbon–phosphorus exchange rate constrains density–speed trade-off in arbuscular mycorrhizal fungal growth. Proceedings of the National Academy of Sciences. 2026;123(6):e2512182123.

18. Birzu G, Hallatschek O, Korolev KS. Fluctuations uncover a distinct class of traveling waves. Proceedings of the National Academy of Sciences. 2018;115(16):E3645–54.

19. Edelstein L. The propagation of fungal colonies: a model for tissue growth. Journal of Theoretical Biology. 1982;98(4):679–701.

20. Davidson F, Sleeman B, Crawford J. Travelling waves in a reaction-diffusion system modelling fungal mycelia. IMA journal of applied mathematics. 1997;58(3):237–57.

21. Held M, Kăspar O, Edwards C, Nicolau DV. Intracellular mechanisms of fungal space searching in microenvironments. Proceedings of the National Academy of Sciences. 2019;116(27):13543–52.

22. Hallatschek O. The noisy edge of traveling waves. Proceedings of the National Academy of Sciences. 2011;108(5):1783–7.

23. Palmer BJ, Johnson CG, Almgren AS, Myers AT, Cannon WR. Three-dimensional modeling of hyphal fusion, branching, and nutrient transport in filamentous fungi. Physical Review E. 2025;112(3):034401.

24. Palmer BJ, Almgren AS, Johnson CG, Myers AT, Cannon WR. BMX: Biological modelling and interface exchange. Scientific Reports. 2023;13(1):12235.

25. Zhang W, Almgren A, Beckner V, Bell J, Blaschke J, Chan C, et al. AMReX: a framework for block-structured adaptive mesh refinement. The Journal of Open Source Software. 2019;4(37):1370.

26. Borgonovo E, Pangallo M, Rivkin J, Rizzo L, Siggelkow N. Sensitivity analysis of agent-based models: a new protocol. Computational and Mathematical Organization Theory. 2022;28(1):52–94.

27. Karniadakis GE, Kevrekidis IG, Lu L, Perdikaris P, Wang S, Yang L. Physics-informed machine learning. Nature Reviews Physics. 2021;3(6):422–40.

28. Lamperti F, Roventini A, Sani A. Agent-based model calibration using machine learning surrogates. Journal of Economic Dynamics and Control. 2018;90:366–89.

29. Wu X, Mcdermott M, Maclean AL. Data-driven model discovery and model selection for noisy biological systems. PLOS Computational Biology. 2025;21(1):e1012762.

30. Lötstedt P. Derivation of continuum models from discrete models of mechanical forces in cell populations. Journal of Mathematical Biology. 2021;83(6):75.

31. Bergman DR, Jackson T, Jain HV, Norton KA. An efficient and flexible framework for inferring global sensitivity of agent-based model parameters. PLoS computational biology. 2025;21(9):e1013427.

32. Brunet E, Derrida B. Shift in the velocity of a front due to a cutoff. Physical Review E. 1997;56(3):2597.

33. Panja D. Effects of fluctuations on propagating fronts. Physics Reports. 2004;393(2):87–174.

34. Brunet E, Derrida B, Mueller AH, Munier S. Phenomenological theory giving the full statistics of the position of fluctuating pulled fronts. Physical Review E—Statistical, Nonlinear, and Soft Matter Physics. 2006;73(5):056126.

35. Vrugt JA, Robinson BA. Improved evolutionary optimization from genetically adaptive multimethod search. Proceedings of the National Academy of Sciences. 2007;104(3):708–11.

36. Bottone EJ, Nagarsheth N, Chiu K. Evidence of self-inhibition by filamentous fungi accounts for unidirectional hyphal growth in colonies. Canadian journal of microbiology. 1998;44(4):390–3.

37. Gutiérrez-Medina B, Vázquez-Villa A. Mechanical interaction between hyphae during three-dimensional growth. Fungal Genetics and Biology. 2021;151:103563.

38. Trinci A. A study of the kinetics of hyphal extension and branch initiation of fungal mycelia. Microbiology. 1974;81(1):225–36.

39. Prosser JI, Trinci A. A model for hyphal growth and branching. Microbiology. 1979;111(1):153–64.

40. Camenzind T, Aguilar-Trigueros CA, Hempel S, Lehmann A, Bielcik M, Andrade-Linares DR, et al. Towards establishing a fungal economics spectrum in soil saprobic fungi. Nature Communications. 2024;15(1):3321.

41. Renardy M, Joslyn LR, Millar JA, Kirschner DE. To Sobol or not to Sobol? The effects of sampling schemes in systems biology applications. Mathematical biosciences. 2021;337:108593.

42. Kolmogorov AN. A study of the equation of diffusion with increase in the quantity of matter, and its application to a biological problem. Moscow University Bulletin of Mathematics. 1937;1:1–25.

43. Van Saarloos W. Front propagation into unstable states. Physics reports. 2003;386(2-6):29–222.

44. Herman J, Usher W. SALib: An open-source Python library for Sensitivity Analysis. The Journal of Open Source Software. 2017 jan;2(9). Available from: https://doi.org/10.21105/joss.00097. doi:10.21105/joss.00097.

45. Saltelli A. Making best use of model evaluations to compute sensitivity indices. Computer physics communications. 2002;145(2):280–97.

46. Pianosi F, Wagener T. A simple and efficient method for global sensitivity analysis based on cumulative distribution functions. Environmental Modelling & Software. 2015;67:1–11.

47. Brown KS, Sethna JP. Statistical mechanical approaches to models with many poorly known parameters. Physical review E. 2003;68(2):021904.

48. Lehmann A, Zheng W, Soutschek K, Roy J, Yurkov AM, Rillig MC. Tradeoffs in hyphal traits determine mycelium architecture in saprobic fungi. Scientific Reports. 2019;9(1):14152.

49. Nardini JT, Baker RE, Simpson MJ, Flores KB. Learning differential equation models from stochastic agent-based model simulations. Journal of the Royal Society Interface. 2021;18(176):20200987.

50. Serghi EU, Kokkoris V, Cornell C, Dettman J, Stefani F, Corradi N. Homo-and dikaryons of the arbuscular mycorrhizal fungus Rhizophagus irregularis differ in life history strategy. Frontiers in Plant Science. 2021;12:715377.

51. Stokes BM, Rogers T, James R. Speed and Shape of Population Fronts with Density-Dependent Diffusion: BM Stokes et al. Bulletin of Mathematical Biology. 2024;86(12):147.

52. Tang J, Riley WJ, Zhu Q. Supporting hierarchical soil biogeochemical modeling: version 2 of the Biogeochemical Transport and Reaction model (BeTR-v2). Geoscientific Model Development. 2022;15(4):1619–32.

53. Sulman BN, Moore JA, Abramoff R, Averill C, Kivlin S, Georgiou K, et al. Multiple models and experiments underscore large uncertainty in soil carbon dynamics. Biogeochemistry. 2018;141(2):109–23.

54. Riley WJ, Sierra CA, Tang J, Bouskill NJ, Zhu Q, Abramoff RZ. Next-Generation Soil Biogeochemistry model representations: A proposed community open-source model Farm (BeTR-S). Multi-scale biogeochemical processes in soil ecosystems: Critical reactions and resilience to climate changes. 2022:233–57.

55. Tang JY, Riley WJ. SUPECA kinetics for scaling redox reactions in networks of mixed substrates and consumers and an example application to aerobic soil respiration. Geoscientific Model Development. 2017;10(9):3277–95.

56. Maherali H, Klironomos JN. Influence of phylogeny on fungal community assembly and ecosystem functioning. science. 2007;316(5832):1746–8.

57. Crowther TW, Van den Hoogen J, Wan J, Mayes MA, Keiser A, Mo L, et al. The global soil community and its influence on biogeochemistry. Science. 2019;365(6455):eaav0550.

